# Lung-selective Cas13d-based nanotherapy inhibits lethal SARS-CoV-2 infection by targeting host protease *Ctsl*

**DOI:** 10.1101/2021.10.03.462915

**Authors:** Zhifen Cui, Cong Zeng, Furong Huang, Fuwen Yuan, Jingyue Yan, Yue Zhao, Jiaoti Huang, Herman F. Staats, Jeffrey I. Everitt, Gregory D. Sempowski, Hongyan Wang, Yizhou Dong, Shan-Lu Liu, Qianben Wang

## Abstract

The COVID-19 pandemic persists as a global health crisis for which curative treatment has been elusive. Development of effective and safe anti-SARS-CoV-2 therapies remains an urgent need. SARS-CoV-2 entry into cells requires specific host proteases including TMPRSS2 and Cathepsin L (Ctsl)^1–3^, but there has been no reported success in inhibiting host proteases for treatment of SARS-CoV-2 pathogenesis *in vivo*. Here we have developed a lung *Ctsl* mRNA-targeted, CRISPR/Cas13d-based nanoparticle therapy to curb fatal SARS-CoV-2 infection in a mouse model. We show that this nanotherapy can decrease lung Ctsl expression in normal mice efficiently, specifically, and safely. Importantly, this lung-selective *Ctsl*-targeted nanotherapy significantly extended the survival of lethally SARS-CoV-2 infected mice by decreasing lung virus burden, reducing expression of pro-inflammatory cytokines/chemokines, and diminishing the severity of pulmonary interstitial inflammation. Additional *in vitro* analyses demonstrated that Cas13d-mediated *Ctsl* knockdown inhibited infection mediated by the spike protein of SARS-CoV-1, SARS-CoV-2, and more importantly, the authentic SARS-CoV-2 B.1.617.2 Delta variant, regardless of TMPRSS2 expression status. Our results demonstrate the efficacy and safety of a lung-selective, *Ctsl*-targeted nanotherapy against infection by SARS-CoV-2 and likely other emerging coronaviruses, forming a basis for investigation of this approach in clinical trials.

## Main

Despite administration of vaccines and changes in human behavior, the coronavirus disease 2019 (COVID-19) pandemic is far from over. The prolonged outbreak demands development of effective therapeutics against severe acute respiratory syndrome coronavirus 2 (SARS-CoV-2), the causative agent of COVID-19. The Food and Drug Administration (FDA) has approved Remdesivir, a nucleoside analog, as a direct-acting antiviral drug against SARS-CoV-2. While Remdesivir shortens time to recovery, it does not provide a survival benefit for COVID-19 patients^4,5^. The FDA has also issued emergency use authorization (EUA) for several monoclonal antibodies, including bamlanivimab plus etesevimab, casirivimab plus imdevimab, and sotrovimab. Clinical studies showed that these monoclonal antibodies can reduce hospitalization and death; however, the effect of monoclonal antibodies is lost or decreased by the emergence of resistant mutations in SARS-CoV-2 variants of concern, especially the B.1.617.2 Delta variant, as well as by unknown bioavailability in affected tissues (e.g. lungs)^6–8^. Thus, development of effective and safe anti-SARS-CoV-2 therapies remains an urgent need.

Recently, virologic and proteomic studies have revealed how SARS-CoV-2 hijacks the host cell machinery during infection^1,9^, which offers knowledge to develop host-directed therapies. Importantly, genome-wide CRISPR/Cas9 screening for host factors has identified the host protease cathepsin L (*CTSL*) as a top-ranked gene not only for SARS-CoV-2 infection, but for related SARS-CoV-1 and Middle East Respiratory Syndrome coronavirus (MERS-CoV) infection in some cell models^2,3^. Indeed, CTSL has long been recognized as an important endosomal cysteine protease that mediates the spike S protein (S) priming and virus entry through virus-host cell endosome membrane fusion^10^. Several CTSL inhibitors have been reported to block entry of coronaviruses (e.g., SARS-CoV-2 and SARS-CoV-1) *in vitro*^1,11–14^ and pseudotyped SARS-CoV-2 infection *in vivo*^15^. However, the antiviral activities of these CTSL inhibitors are lost or significantly reduced upon expression of the serine proteases for cell entry such as TMPRSS2 on the plasma membrane^1,16^. Moreover, a potent CTSL inhibitor against SARS-CoV-1 infection *in vitro* has failed to provide survival benefit in a lethal SARS-CoV-1 mouse model^13^. It remains unclear whether the lack of antiviral potency of CTSL inhibitors *in vivo* and in some *in vitro* models is due to the use of alternative coronavirus entry pathways and issues of drug efficacy or delivery. It is currently unknown whether targeting CTSL would block authentic SARS-CoV-2 infection *in vivo*. In this study, we have developed a lung *Ctsl* mRNA-targeted, CRISPR/Cas13d-based lipid nanoparticle (LNP) therapy to inhibit fatal SARS-CoV-2 infection in mouse models, as well as SARS-CoV-1, SARS-CoV-2, and its Delta variant infection in TMPRSS2-expressing cultured cells.

## Development of a lung-selective Cas13d-based nanotherapy targeting *Ctsl* mRNA

As the primary organ of SARS-CoV-2 infection is the lung, we first used selective organ targeting (SORT) nanotechnology^17^ to generate lung-selective LNP by adding the cationic lipids DOTAP at 50% molar percentages to the traditional LNP formulation employed by the FDA-approved RNAi therapy Patisiran/Onpattro^18^ (Fig. 1a). The selective lung targeting effects were visualized by bioluminescence imaging of both animal whole body and major organs 3 h following intravenous (IV) administration of characterized LNPs encapsulating a luciferase mRNA (LNP-Luc) into normal mice (Fig. 1b, Extended Data Fig. 1a-d). We next constructed *Ctsl* mRNA-targeting LNPs by using the lung-selective LNPs to encapsulate CRISPR/CasRx (a Cas13d RNA-targeting enzyme from the *Ruminococcus flavefaciens* strain XPD3002)^19^ and an unprocessed guide RNA (pre-gRNA) recognizing mouse *Ctsl* mRNA (Extended Data Fig. 2a, b). Since previous studies found that co-delivery of *Cas9* mRNA with gRNAs into cells produces faster gene editing kinetics and fewer off-target effects compared with a plasmid encoding them^20,21^, we engineered lung-targeting LNPs to deliver *CasRx* mRNA and pre-gRNA oligos targeting *Ctsl* in lungs (called LNP-*CasRx*-pre-g*Ctsl*). *CasRx* mRNA expression in lungs peaked at 4 h after LNP-*CasRx* injection and then markedly decreased at 24 h (Extended Data Fig. 2c), similar to the fast and transient kinetics of *Cas9* mRNA^22^. Compared with three control groups, including DPBS, empty LNPs (eLNPs), and LNP-*CasRx*-pre-gControl, IV administration of LNP-*CasRx*-pre-g*Ctsl* significantly inhibited mRNA and protein expression of Ctsl in mouse lungs (Fig. 1c, d). These findings were further validated by immunohistochemistry (IHC) of lung sections. The strong Ctsl staining of bronchiolar epithelial cells, type II alveolar epithelial cells and macrophages was markedly decreased by LNP-*CasRx*-pre-g*Ctsl* treatment (Fig.1e). Notably, *Ctsl* expression was not changed in liver or spleen, and expression of other cathepsin family members *Ctsd, Ctsb, Ctss* was not altered in lungs after LNP-CasRx-pre-g*Ctsl* treatment (Extended Data Fig. 2d, e), demonstrating the tissue and gene specificity of *Ctsl* targeting by LNP-*CasRx*-pre-g*Ctsl*. Furthermore, no significant changes in cytokine/chemokine expression were observed in lungs following LNP-*CasRx*-pre-g*Ctsl* treatment (Extended Data Fig. 3a). Finally, mouse liver or kidney function and hematology (including red blood cell count, hemoglobin level, white blood cell count and platelet count) were normal (Fig. 1f), with no histological toxicity in major organ tissues of LNP-*CasRx*-pre-g*Ctsl* treated mice (Extended Data Fig. 3b). Taken together, these data demonstrate that the lung-selective LNP-*CasRx*-pre-g*Ctsl* system can decrease lung Ctsl expression efficiently, specifically, and safely.

**Fig. 1.**
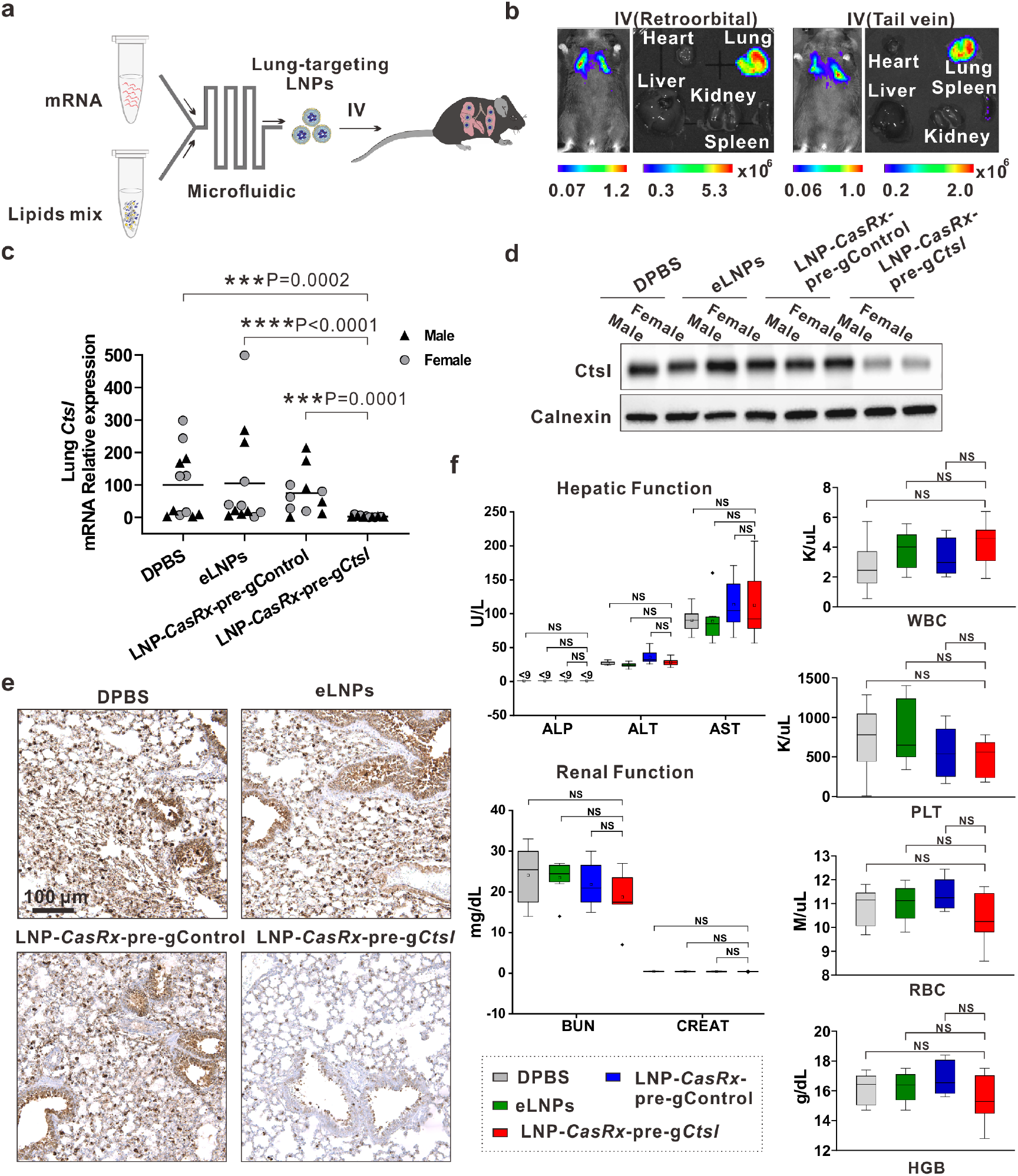
Ctsl is specifically knocked down by a lung-selective LNP encapsulating *CasRx*-pre-gRNA system in normal C57BL/6 mice with no toxicity. **a,** Schematic description for the generation of a lung-targeting LNP encapsulating RNAs and its administration by IV injection. **b,** IVIS imaging for the LNP biodistribution. Animal’s whole body, and organs were imaged 3 h after IV injection (retroorbital or tail vein) of LNPs loaded with luciferase mRNA at 0.5 mg/kg. **c-f,** Ctsl mRNA and protein levels are significantly reduced in mouse lung with minimal systemic toxicity. Mice were given DPBS, empty LNPs, LNP-*CasRx*-pre-gControl or LNP-*CasRx*-pre-g*Ctsl*, respectively, by tail vein. After 72 h, samples were collected for analysis. **c,** *Ctsl* mRNA level in lung. *Ctsl* transcript levels are normalized by *Gapdh* and presented as the percentage of the DPBS group. Each group consists of 6 females and 6 males except 5 females for LNP-*CasRx*-pre-gControl group. Bar shows the average, and each label represents an individual mouse. *P* values were calculated by one-tailed Mann-Whitney u test, grand mean. *** *P*<0.001, \*\*\*\**P*<0.0001. **d,** Representative western blot of Ctsl protein in lung. One male and one female mouse were selected for each group. Calnexin was used as a loading control. The Ctsl mature form was shown. **e,** Representative images for Ctsl immunostaining in lung. One section from 1 mouse from each control group and 3 mice from LNP-*CasRx*-pre-g*Ctsl* group were subjected to IHC analysis. **f,** Hepatic and renal functions as well as hematology are not impaired by LNP-*CasRx*-pre-*Ctsl* treatment. ALP, alkaline phosphatase; ALT, alanine transaminase; AST, aspartate aminotransferase; BUN, blood urea nitrogen; CREAT, Creatinine; WBC, white blood cells; PLT, platelets; RBC, red blood cells; HGB, hemoglobin. Data are presented as box and whiskers of each group with n=8 (4 females and 4 males in each group except 3 females and 5 males in LNP-*CasRx*-pre-gControl group). Whiskers are min to max. *P* values were calculated by two-tailed Mann-Whitney u test, NS, not significant.

## LNP-*CasRx*-pre-g*Ctsl* treatment extended survival of lethally SARS-CoV-2-infected mouse

The K18-hACE2 mouse model establishes a robust SARS-CoV-2 infection and recapitulates major features of human severe COVID-19, such as severe interstitial pneumonia and exuberant inflammatory response^23–28^. We thus chose this lethal challenge model to evaluate the antiviral efficacy of our nanotherapy *in vivo* (Fig. 2a). When challenged with 10^5^ plaque-forming unit (PFU) of SARS-CoV-2 (USA-WA1/2020), the mice in the three control groups displayed a progressive weight loss and symptom manifestation until they all succumbed within 8 days after inoculation (Fig. 2b-d). By contrast, mice administered with LNP-*CasRx*-pre-g*Ctsl* two days prior to and one day post infection (DPI) with SARS-CoV-2 exhibited a delayed onset and reduced the extent of the weight loss and clinical symptoms (Fig. 2b, c), contributing to the significantly increased survival (Fig. 2d).

**Fig. 2.**
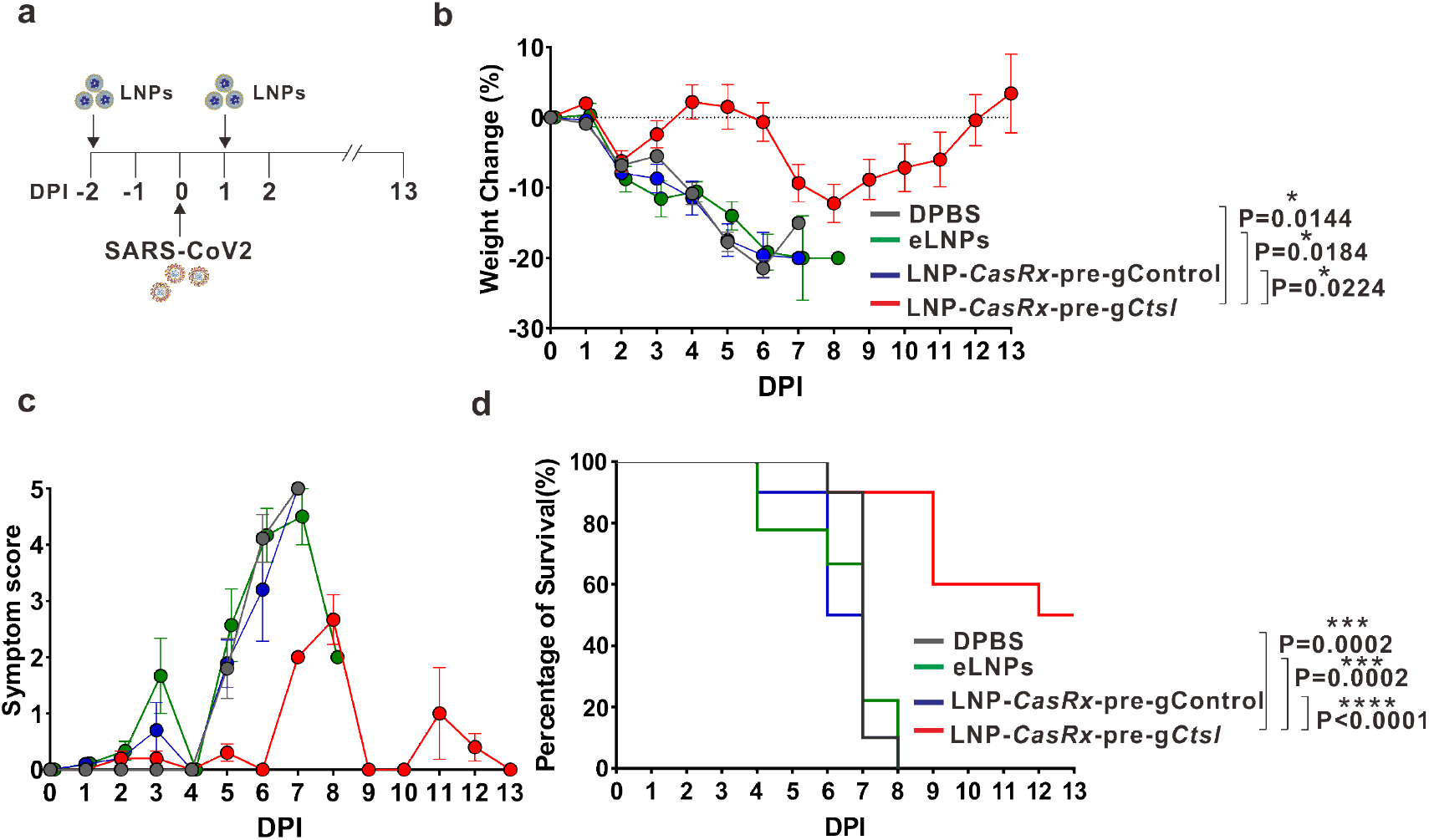
LNP-*CasRx*-pre-g*Ctsl* treatment protects K18-hACE2 mice against lethal SARS-CoV-2 infection. **a,** Schematic illustration of the experiment design. Seven-to eight-week-old K18-hACE2 female and male mice were intranasally infected with 10^5^ PFU of SARS-CoV-2 (USA-WA1/2020). They were treated with DPBS, eLNPs, LNP-*CasRx*-pre-gControl or LNP-*CasRx*-pre-g*Ctsl* through the retroorbital injections 2 days prior to and 1 day post infection (DPI). Each group consists of 10 mice (5 females and 5 males) except 9 for eLNPs group (5 females and 4 males). Mice weights and clinical signs were monitored daily to DPI 13. **b,** Percentage of weight change. Data are presented as mean ± SEM. The dotted line represents the initial body weight. Mice that loss >20% of their initial body weight were humanely euthanized. *P* values were calculated by two-tailed *Student*’s *t*-tests, * *P*<0.05. **c,** Symptom score. Each animal was evaluated daily based on the criteria: 0, normal; 1, ruffled fur and hunched; 2, respiration-labored breathing and sneezing; 3, discharge from nose; 4, lethargic; 5, moribund or dead. Data are presented as mean ± SEM. **d,** Kaplan-Meyer survival curves. *P* values were determined by logrank (Mantel-Cox) test. *** *P*<0.001, **** *P*<0.0001.

## LNP-*CasRx*-pre-g*Ctsl* treatment reduced lung viral burden

With an inoculation of 10^5^ PFU of SARS-CoV-2, viral burden in K18-hACE2 mouse lungs has been shown to peak at 2 DPI^23,25^. We thus collected the lung tissues at this time point to evaluate the efficacy of LNP-*CasRx*-pre-g*Ctsl* treatment in reducing the lung viral load (Fig. 3a). Since there were no differences observed among three control groups (DPBS, eLNPs and LNP-*CasRx*-pre-gControl) for mouse survival, weight loss and clinical signs (Fig. 2b-d), we pooled them as one “Control” group for all the following analysis. We found that LNP-*CasRx*-pre-g*Ctsl* treatment decreased the lung virus infectivity by two orders of magnitude compared with the Control group (Fig. 3b). Consistently, we observed significantly lower mRNA and/or protein expression of viral nucleocapsid (N) and envelope (E) in LNP-*CasRx*-pre-g*Ctsl* treatment group than in the Control group (Fig. 3c, d, h, i). Consistent with previous reports^25,28^, the lung sections of the Control group showed strong diffuse N protein staining at 2 DPI, which was significantly decreased by LNP-*CasRx*-pre-g*Ctsl* treatment (Fig. 3e, f). As expected, LNP-*CasRx*-pre-g*Ctsl* treatment significantly decreased mRNA and protein expression of Ctsl in lungs (Fig. 3g-i). Thus, LNP-*CasRx*-pre-g*Ctsl* treatment ameliorated SARS-CoV-2 infection efficiently and rapidly.

**Fig. 3.**
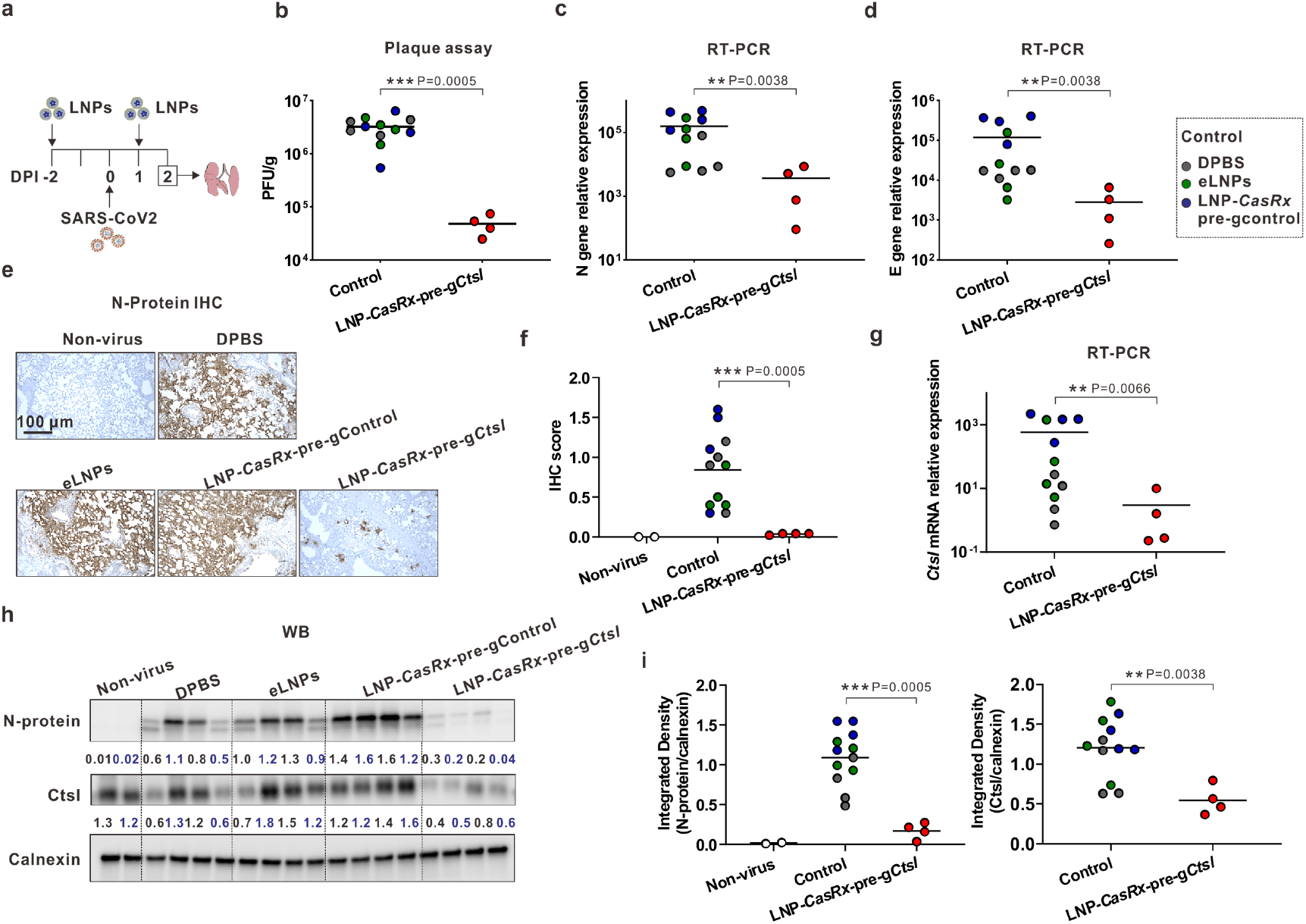
CasRx-mediated Ctsl knockdown markedly decreases SARS-CoV-2 burden in lungs of infected K18-hACE2 mice. **a**, Schematic illustration of the experiment design. Same treatment as in Fig. 2. Each group consists of 2 females and 2 males. Lungs were collected for analysis at 2 DPI. Groups of DPBS, eLNPs, and LNP-*CasRx*-pre-gControl were pooled as Control. **b**, Infectious viral titer in lung. Tissue homogenates were analyzed by plaque assay. **c**, Viral nucleocapsid N gene transcript levels. **d**, Viral envelope E gene transcript levels. **e**, Representative images for viral N protein immunostaining in lung. One section of lung tissues from 4 mice of each group were subjected to IHC analysis. **f**, IHC score for the viral N protein. The score of N protein expression in lung sections is the product of intensity (graded as 0-absent, 1-weak positive, 2-strong positive) and proportion of staining in pneumocytes showing maximum intensity. **g**, *Ctsl* transcript levels. **h**, Western blot of viral N protein and Ctsl mature form protein in lung. Calnexin was used as a loading control. Each lane represents an individual mouse. The ratio of N or Ctsl protein over calnexin is listed under the blot. Non-virus, two mice (1 female and 1 male) were not challenged with virus. **i**, Integrated density of N protein (left panel) and Ctsl (right panel) over the loading control calnexin. **c, d**, **g** all transcript levels are normalized by *Gapdh*. Bar is the average of the group. Each circle represents an individual animal. *P* values were calculated by one-tailed Mann-Whitney u test, grand mean. ** *P*<0.01, *** *P*<0.001.

## LNP-*CasRx*-pre-g*Ctsl* treatment reduced cytokine/chemokine levels and lesions in lungs

During the development of severe COVID-19 disease, uncontrolled inflammation termed a cytokine storm is believed to be responsible for multi-organ damage and fatal coutcome^29,30^. For example, elevated CXCL10 and TNF levels are correlated with increased disease severity and decreased survival of COVID-19 patients^29,31^. Here we showed that a lethal dose of SARS-CoV-2 infection significantly increased the transcript levels of *Cxcl10* and *Tnf* in mouse lung at 4 DPI in the Control group as compared with unchallenged mice (Fig. 4a, b). Remarkably, *Cxcl10* and *Tnf* expression in LNP-*CasRx*-pre-g*Ctsl* treated group was almost the same as that in unchallenged mice (Fig. 4b). Similarly, *Ccl5, Ccl2* and *Isg15* levels were all elevated by virus infection but decreased by LNP-*CasRx*-pre-g*Ctsl* treatment (Extended Data Fig. 4). Histologically, the virus-infected Control group mice had parenchymal lung lesions that were patchy in distribution at 4 DPI. Perivascular lymphoid infiltrates and interstitial thickening of alveolar septal regions with mixed mononuclear and polymorphonuclear leukocyte infiltrations were the most prominent features of the lung lesions, and thus were scored in the treatment and control groups to provide a semi-quantitative pathology index. Notably, this index was decreased in the LNP-*CasRx*-pre-g*Ctsl* group vs. the Control group (Fig. 4c, d). Similar to its effects at 2 DPI, LNP-*CasRx*-pre-g*Ctsl* also reduced the mRNA and/or protein expression level of viral N and E in infected lungs at 4 DPI (Extended Data Fig. 5a-f). The efficient knockdown of Ctsl in mouse lung by LNP-*CasRx*-g*Ctsl* treatment was also confirmed (Extended Data Fig. 5c, d, g). These data indicate that LNP-*CasRx*-pre-g*Ctsl* treatment inhibited the expression of pro-inflammatory cytokines/chemokines and reduced lung disease.

**Fig. 4.**
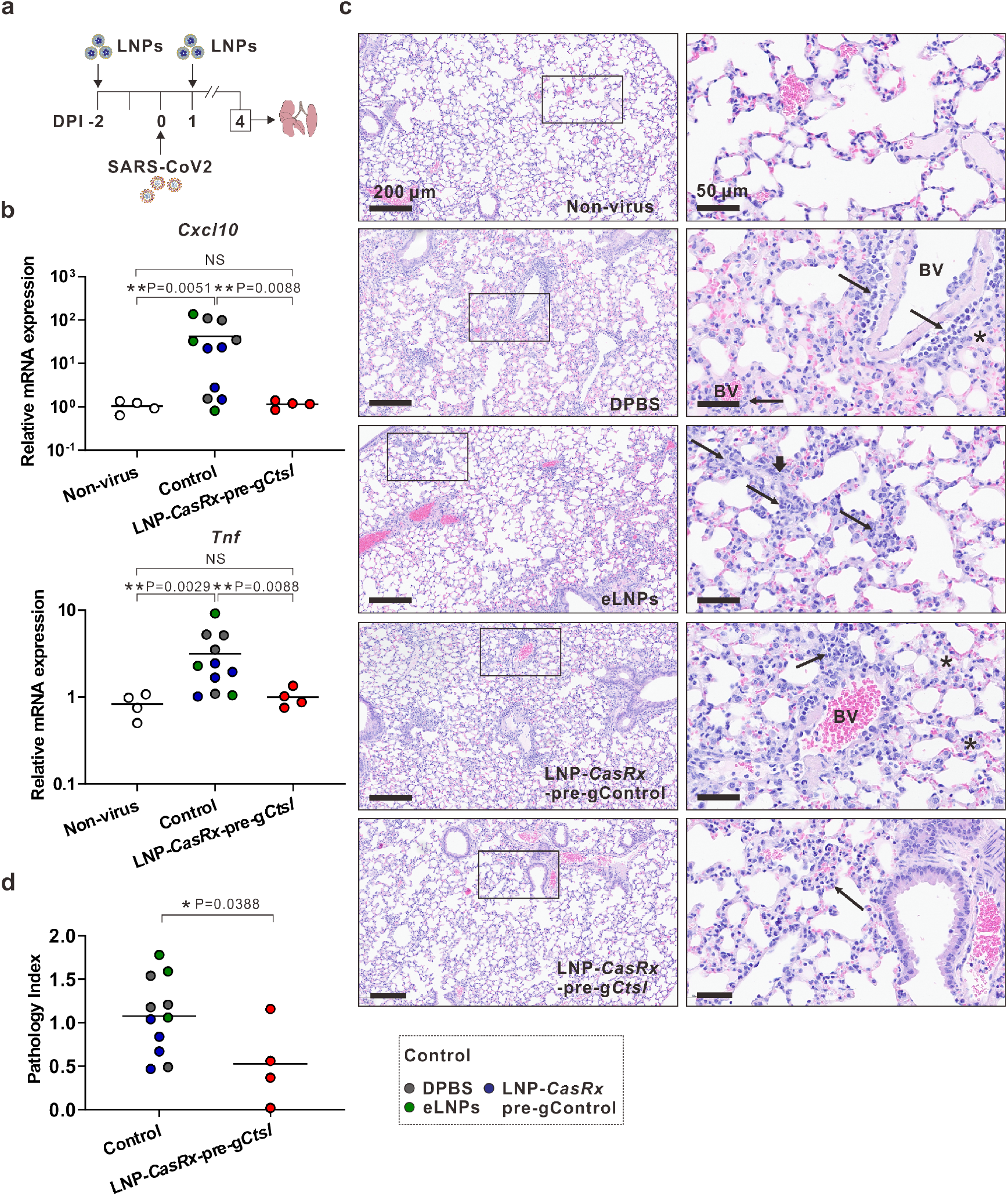
LNP-*CasRx*-pre-g*Ctsl* reduces chemokine/cytokine levels and lung pathogenesis in SARS-CoV-2-infected K18-hACE2 mice. **a**, Schematic illustration of the experiment design. Same treatment as in Fig. 2. Each group consists of 2 females and 2 males except for eLNPs group (1 female and 2 males). Lungs were collected for analysis at 4 DPI. Groups of DPBS, eLNPs, LNP-*CasRx*-pre-gControl were pooled as Control. **b**, Transcript levels of *Cxcl10* and *Tnf* in lung. All transcript levels are normalized by *Gapdh*. **c**, Representative photomicrographs of H&E-stained lung sections demonstrate the perivascular, interstitial and alveolar inflammatory lesions in virus-infected mice were reduced by LNP-*CasRx*-pre-g*Ctsl* treatment. Annotations: BV, blood vessel; thin arrow, perivascular or interstitial inflammatory infiltrates; thick arrow, karyorrhexic nuclei and region of diffuse alveolar damage; asterisk, intra-alveolar edema. **d**, Lung pathology index. Randomly selected regions of interest in each of four animals per group were scored for inflammatory lesions as outlined in the Methods. *P* values were calculated by one-tailed Mann-Whitney u test, grand mean. NS, not significant, * *P*<0.05, \*\**P*<0.01.

While primarily a respiratory disease, COVID-19 patients may suffer neurological complications ranging from loss of smell and headache to acute encephalopathy^32–34^. Here we observed high expression levels of viral N and E genes in 5 out of 11 mice in the Control group (Extended Data Fig. 6a-d), consistent with previous observations that some mice with high dose virus challenge may develop brain infection^25,26,28^. Extensive staining of the viral N protein was mainly located in cerebral cortex, thalamic and hypothalamic regions (Extended Data Fig. 6d). Of note, the viral transcript expression and N protein staining levels were much less in the LNP-*CasRx*-pre-g*Ctsl* group although the statistical significance was not reached (Extended Data Fig. 6b-d), possibly due to the heterogenous response to SARS-CoV-2 infection in the brain. Clinically, some COVID-19 patients also displayed a significant increase in the levels of CCL and CXCL family chemokines in brain, possibly caused by indirectly relayed inflammation into the brain by SARS-CoV-2 infection^35^. We also observed an elevation of the transcripts of *Cxcl10, Ccl2* and *Ccl5*, particularly in some control mice, whereas their expression in LNP-*CasRx*-pre-g*Ctsl* group remained at a similar level as that in unchallenged mice (Extended Data Fig. 6e). Therefore, LNP-*CasRx*-pre-g*Ctsl* treatment likely protects brain from direct and/or indirect damages by SARS-CoV-2 infection. Histological changes in the brain were limited to focal meningeal and perivascular accumulation of inflammatory cells in Virchow-Robin spaces. The limited tissue sampling scheme did not allow comparison of this lesion across the Control and treatment groups (Extended Data Fig. 6f).

Finally, no significant histological changes were present in other tissues examined (liver, spleen, kidney, jejunum, and colon), whether associated with disease or treatments at 4 DPI (Extended Data Fig.7).

## CasRx-mediated *Ctsl* knockdown inhibited authentic B.1.617.2 Delta variant infection *in vitro*

Although previous studies found that antiviral activities of Ctsl inhibitors are lost or markedly decreased in cells expressing TMPRSS2^1,13,16^, we found that *CasRx*-pre-g*Ctsl* delivered by Lipofectamine MessengerMAX (a commercial transfection reagent) similarly inhibited the pseudotyped SARS-CoV-2 infection of TMPRSS2-negative /CTSL-positive Vero-ACE2 cells in the absence or presence of exogenous TMPRSS2 expression (Fig. 5a, b, Extended Data Fig. 8a, b). Additionally, *CasRx*-pre-g*Ctsl* transfection reduced entry of both pseudotyped SARS-CoV-2 and SARS-CoV-1 but not that of vesicular stomatitis virus (VSV) infection in CTSL-positive Caco-2 cells expressing endogenous TMPRSS2 (Fig. 5c). Importantly, CasRx-mediated *Ctsl* knockdown also significantly inhibited infection of authentic SARS-CoV-2 B.1.617.2 Delta variant in Vero-ACE2-TMPRSS2 cells (Fig. 5d, e). Collectively, these data demonstrated that CasRx-mediated *Ctsl* knockdown effectively inhibited infection by SARS-CoV-2 and related coronaviruses, including variants of concern, regardless of the TMPRSS2 expression status.

**Fig. 5.**
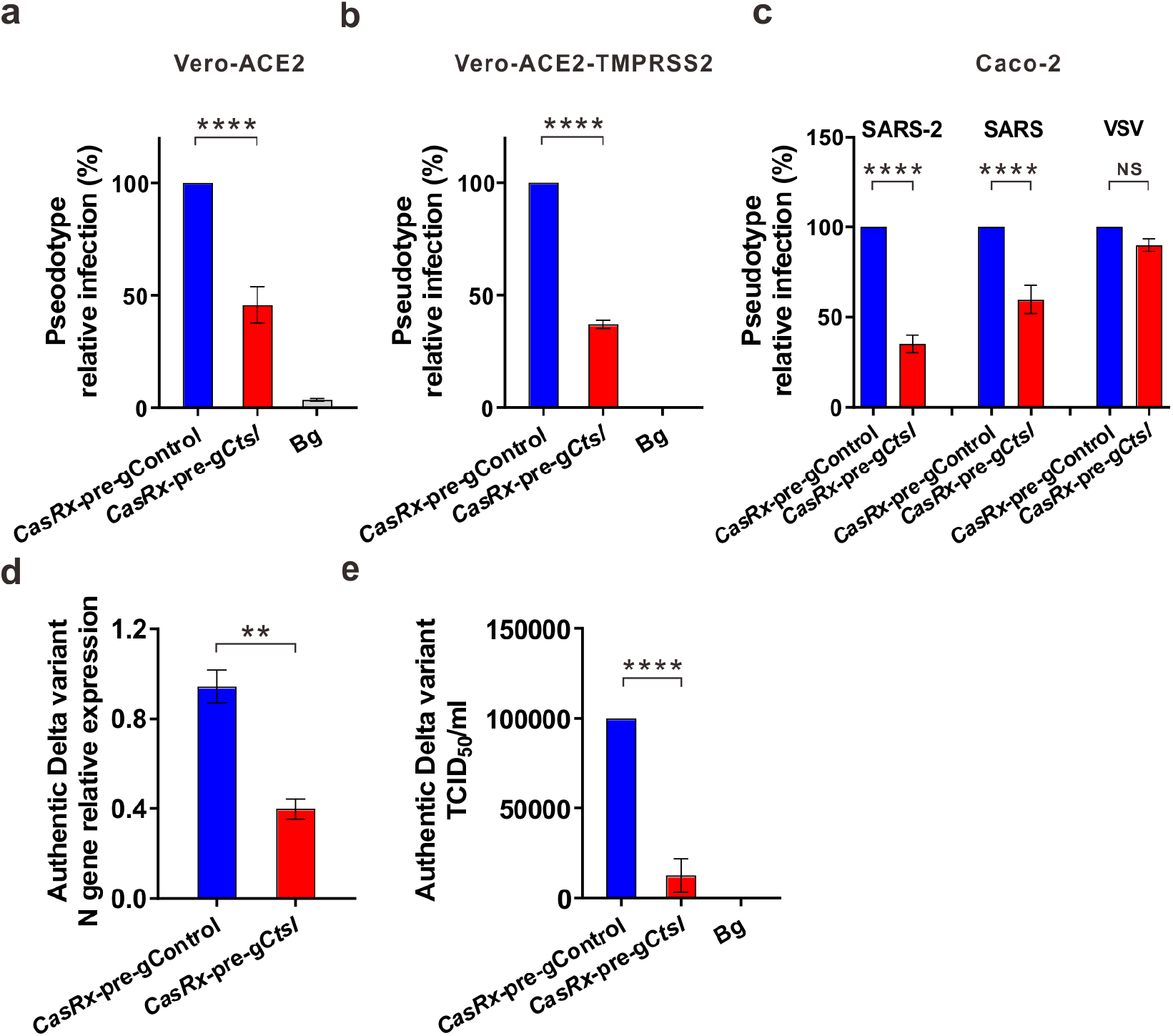
CasRx-mediated *CTSL* knockdown inhibits pseudotyped coronaviruses and authentic SARS-CoV-2 Delta variant (B.1.617.2) infection in cells. **a-c,** Pseudotyped virus relative infection in Vero-ACE2 (**a**), Vero-ACE2-TMPRSS2 (**b**) and Caco-2 (**c**) cells. Cells were pretreated with LNP-*CasRx*-pre-g*Ctsl* for 24 h, followed by incubation with infectivity-normalized pseudovirus SASR-CoV-2 (SARS-2), SARS-CoV-1 (SARS) or VSV. **d**-**e**, *CasRx*-g*CTSL* treatment reduces live SARS-CoV-2 B.1.617.2 Delta variant infection in Vero-ACE2-TMPRSS2 cells. **d**, Viral N gene transcript level as normalized by 18s rRNA. **e**, Virus infectivity as determined by TCID_50_. Bg, background, cells were not challenged with virus and used as negative control. Experiments were performed in triplicate. Data are presented as mean± SEM. *P* values were calculated by two-tailed *Student*’s t-tests. NS, not significant; ** *P*<0.01; **** *P*<0.0001.

## Discussion

Despite the well-recognized role of the host protease CTSL in mediating SARS-CoV-2 cell entry^1–3^, there has been no or limited success using CTSL inhibitors to inhibit live SARS-CoV-2 infection *in vivo*. We propose that gene therapy may be a critical strategy for targeting host proteases with greater efficiency, as it impairs functions of both catalytic and non-catalytic domains of the proteases^36^. Since cathepsins play important physiological roles in many processes such as immune response, autophagy and development^10,37^, we have used the recently developed CRISPR/CasRx RNA editing system^19^ to transiently knockdown *Ctsl* mRNA rather than using CRISPR/Cas9 to permanently delete the *Ctsl* gene. CasRx has been reported to mediate highly efficient and specific RNA knockdown^19^ without the widespread off-target effects that are associated with RNA interference (RNAi) strategies due to their key roles in endogenous processes^38,39^. Indeed, the potent *Ctsl* mRNA knockdown by the CasRx (Fig. 1c) does not affect the expression of other Ctsl family members (Extended Data Fig. 2e) that are involved in critical physiological and pathological processes (e.g., Ctss functions in antigen processing/presentation and antibody production^40,41^). Furthermore, the non-pathogenic *Ruminococcus flavefaciens*-derived CasRx appears to be non-inflammatory and non-toxic *in vivo* (Fig. 1f, Extended Data Fig. 3). Finally, the delivery of *CasRx/gCtsl* by lung-targeting LNPs allows a selective knockdown of Ctsl in lungs (Fig. 1a-e, Extended Data Fig. 2d) without affecting the Ctsl expression in other organs where it may have important roles (e.g., Ctsl inhibition may impair CD4+ T cell selection in splenocytes and thymocytes^42,43^). Collectively, our newly developed lung *Ctsl*-targeted nanotherapy strategy contributes to an efficient, specific, and safe Ctsl knockdown in lungs. Importantly, LNP-*CasRx*-pre-g*Ctsl* treatment significantly protects K18-hACE2 transgenic mice from lethal infection by SARS-CoV-2 (Fig. 2). The survival benefit of this nanotherapy is believed to be contributed by reduction of viral load, cytokines/chemokines levels, and lung pathology (Fig. 3, 4). Interestingly, our *in vitro* analysis found that CasRx-mediated *Ctsl* knockdown can block SARS-CoV-2 infection of both TMPRSS2-positive and -negative cells (Fig. 5a-c), highlighting an indispensable role of CTSL in multiple SARS-CoV-2 entry pathways. We also examine the effect of CasRx-mediated *Ctsl* knockdown on infection of the authentic B.1.617.2 Delta variant, which is currently the predominant SARS-CoV-2 variant in the United States. Compared with other variants, the Delta variant is more contagious, which is at least in part caused by markedly increased cell entry due to enhanced spike protein cleavage^8^. Importantly, CasRx-mediated *Ctsl* knockdown significantly inhibits live B.1.617.2 Delta variant infection of TMPRSS2-expressing cells (Fig. 5d, e), suggesting that Ctsl may play an essential role in cleaving Delta variant spike protein and increase its virulence.

SARS-CoV-2 is one of three novel coronaviruses (also including SARS-CoV-1 and MERS-CoV) that have been found to cause large-scale pandemics in the 21^st^ century^44^. Importantly, our LNP-*CasRx*-pre-g*Ctsl* treatment also inhibits SARS-CoV-1 pseudovirus infection in TMPRSS2-expressing cells (Fig. 5c), suggesting that lung *Ctsl*-targeted nanotherapy may also inhibit infection of other related highly pathogenic coronaviruses *in vivo*. In addition, the findings that many coronaviruses circulating in bats capable of replicating in human lung and airway cells necessitate the need for the development of pan antiviral therapeutic strategies for future zoonotic coronavirus outbreaks^44^. Given that CTSL is involved in mediating cell entry of several coronaviruses as well as their mutants^2,3^ (Fig. 5), and considering the fact that the vast majority of coronavirus spike do not undergo frequent mutations in the CTSL cleavage sites (between S1/S2 border and the S2’ position^14,15^), our lung *Ctsl*-targeted nanotherapy may represent an attractive therapeutic approach. It is also noteworthy that the RNA-editing system allows the flexible design of gRNAs in *CasRx*-based LNPs that can target any other essential host factors related to coronavirus infection^2,3^.

## Methods

### Preparation of lung-targeting lipid nanoparticles encapsulating *CasRx* mRNA and gRNA oligos

Lipid nanoparticles (LNPs) were generated by simultaneously mixing one volume of lipid mixture (25 : 5: 19.3 : 0.8 : 50 molar ratio) of DLin-MC3-DMA (MedKoo Biosciences, #555308), DSPC (1,2-distearoyl-sn-glycero-3-phosphocholine, Avanti Polar Lipids, #850365), Cholesterol (Sigma, #C3045), DMG-PEG2000 (NOF America Corporation), and DOTAP (1,2-dioleoyl-3-trimethylammonium-propane, Avanti Polar Lipids, #890890) in ethanol and two volumes of RNAs in citrate buffer (10 mM, pH 4.0) in a microfluidic device (TheNanoAssemblr® SparkTM, Precision Nanosystems, BC, Canada) to satisfy a final weight ratio of 20 :1 for total lipids : RNAs. *CasRx* mRNA with pseudouridine modification (pseudouridine-5’-triphosphate, TriLink) was synthesized using *in vitro* transcription (AmpliScribe T7-Flash Transcription Kit, Lucigen) and was then installed with 5’cap (Vaccinia Capping System, NEB, USA; Cap 2’-O-methyltransferase, NEB, USA) and 3’ Poly(A) tail structures by Dr. Yizhou Dong’s lab at The Ohio State University^45,46^. Pre-gControl and pre-g*Ctsl* oligos with 2’OMe and phosphorothioate modification were synthesized by Exonano RNA, LLC (Columbus, Ohio), and their sequences are listed in Extended Data Table 1. The formulation was then dialyzed using Slide-A-Lyzer® Dialysis cassette (WMCO 3.5 kDa, Thermo Fisher, #66330) against 1 X DPBS for 1 h, and diluted to 2 μg/μl of lipids with DPBS for injections.

### Physicochemical characterization of LNPs

Empty LNPs (eLNPs) and luciferase mRNA (TriLink, #L-7202)-loaded LNPs (LNP-Luc mRNA) were characterized by their size, morphology, zeta potential and stability. Size and zeta potential were measured using capillary cells (Malvern, #DTS1070) by Zetasizer Nano ZS (Malvern Panalytical). Transmission Electron Microscope (TEM) examination was performed by Chapel Hill Analytical and Nanofabrication Laboratory, University of North Carolina-Chapel Hill. Briefly, LNPs were stained with 2% uranyl acetate for 2 mins and imaged by Talos F200X (Thermo Fisher) at an accelerating voltage of 200 kV. We also stored the LNPs in DPBS at 4°C for ten days to study their stability. Aliquots were taken every other day for determination of their encapsulation efficiency by RiboGreen assay (Thermo Fisher, #R11490).

### Mouse experiments and authentic SARS-CoV-2 virus infection of mice

Animal studies were approved by the Institutional Animal Care and Use Committee of Duke University. C57BL/6 mice (Stock# B6-MPF, 5-8 weeks old, female or male) were purchased from Taconic (Rensselaer, NY). Transgenic mice expressing human ACE2 under control of cytokeratin 18 promoter K18-hACE2 (Stock# 034860, 7-8 weeks old, female or male) were obtained from Jackson Laboratory (Bar Harbor, ME). Mice were maintained on a 12 h light/dark cycle with food and water ad libitum and acclimatized for a week before the beginning of an experiment. All animal experiments with SARS-CoV-2 were performed in the Duke Regional Biocontainment Laboratory, a Biosafety Level 3 (BSL3) facility.

#### IVIS imaging

LNPs encapsulating luciferase mRNA (TriLink, #L-7202) were generated as described above and given to C57BL/6 mice by IV bolus through either retroorbital route (n=4) or tail vein (n=4) at a dose of 0.5 mg/kg mRNA. Three h later, mice were intraperitoneally injected with the substrate D-Luciferin (150 mg/kg, Perkin Elmer, #770505). Whole body and major organs (heart, liver, spleen, lung, kidney) were then imaged for fluorescence using an IVIS Lumina XR system (Perkin Elmer).

#### Toxicity profiling

C57BL/6 mice were used for *in vivo* toxicity studies. Mice were injected through tail vein with DPBS, empty LNP, LNP-*CasRx*-pre-gControl or LNP-*CasRx*-pre-g*Ctsl*, respectively. Each group had 8 mice with 4 female and 4 males. Blood was collected 72 h after the injection. Complete blood cell count (CBC) and serum level of alkaline phosphatase (ALP), alanine aminotransferase (ALT), aspartate aminotransferase (AST), blood urea nitrogen (BUN) and creatinine (CREAT) were assessed by the Animal Histopathology and Laboratory Medicine Core, University of North Carolina-Chapel Hill. Selected tissues, including lung, liver, spleen, heart, and kidney, were collected for histopathological evaluation.

#### Survival study

K18-hACE2 mice were lightly anesthetized with isoflurane and infected intranasally with 10^5^ PFU of SARS-CoV-2 (USA-WA1/2020 strain) in a total volume of 50 μl DMEM on Day 0. Mice were given with DPBS (n=10), empty LNP (n=9), LNP-*CasRx*-pre-gControl (n=10) or LNP-*CasRx*-pre-g*Ctsl* (n=10), respectively, through retroorbital route 2 days prior to infection and 1 day post infection (DPI). Animal weight, temperature, and clinical signs were monitored daily. The criteria for the symptom scoring are: 0, normal; 1, ruffled fur and hunched; 2, respiration-labored breathing and sneezing; 3, discharge from nose; 4, lethargic; 5, moribund or dead. Mice were humanely euthanized when weight loss >20% and/or symptom score reaches 5.

#### Sample collection

Organ tissues were collected on 2 or 4 DPI as indicated. Left lung lobe, left brain, liver, spleen, kidney, jejunum and colon were immersion fixed in 10% neutral buffered formalin (NBF) for histological analysis. Right lung lobe and right brain were homogenized and used for extraction of total RNA and proteins and plaque assay.

### Histology and immunohistochemistry

Tissues were processed routinely in paraffin, sectioned at four microns, stained with hematoxylin & eosin (H&E), and evaluated by a board-certified veterinary pathologist in a masked manner without knowledge of study allocation group. A pathology disease index of respiratory disease was created using semi-quantitative scoring on digitized lung sections (Aperio AT2, Leica Biosystems) with Imagescope (Leica Biosystems) software. Scoring consisted of assigning a 0-4 severity score (0-normal – 4 most severe) for the two most consistent pathology parameters (perivascular lymphoid cuffing and interstitial alveolar thickening with inflammatory cells) in the model. A pathology index score was made by multiplying the severity score of each parameter by the area of the lung involved and combining the two parameter scores. The percentage of pulmonary parenchyma affected was assessed on three randomly selected regions of interest from the digitized whole slide images using a grid overlay with Imagescope software. Immunohistochemistry (IHC) was performed by the Research Immunohistology Shared Resource at Duke University. Antibodies targeting the CTSL (R&D SYSTEMS, #AF1515, 1:800) and the SARS-CoV-2 nucleoprotein (N) (Sino Scientific, #40143-R019, 1:20000) were used. The IHC score of nucleocapsid protein expression in lung sections ranged from 0 to 2, which was performed by the product of intensity (graded as 0-absent; 1-weak positive; 2-strong positive) and proportion of staining in pneumocytes showing maximum staining intensity.

### Cell culture

HEK293T (ATCC #CRL-11268, RRID: CVCL_1926), Vero-ACE2 (Vero-E6 expressing high endogenous ACE2, BEI, NR-53726) cells were grown in Dulbecco’s modified Eagle’s medium (DMEM) supplemented with 1% penicillin/streptomycin and 10% fetal bovine serum (FBS, Thermo). Caco-2 (ATCC #HTB-37, RRID: CVCL_0025) cells were grown in DMEM supplemented with 1% penicillin/streptomycin and 20% FBS. Vero-ACE2 cells stably expressing TMPRSS2 were generated by transduction of Vero-ACE2 cells with a lentiviral vector expressing human TMPRSS2, followed by blasticidin S HCl selection (7.5 μg/mL) for 7 days. All cell lines were maintained at 37°C with 5% CO_2_. The cell lines were routinely screened using VenorTM GeM Mycoplasma Detection Kit (Sigma-Aldrich, #MP0025) following the manufacture’s protocol.

### RNA isolation and qRT-PCR

Total RNA of cells was isolated by RNeasy mini kit (Qiagen, #74104). Tissues were first homogenized by a BeadBug microtube homogenizer (for virus-infected tissues) or a power homogenizer (for normal tissues), total RNA was then extracted using Ambion® PARIS™ system (Invitrogen, #AM192) and reverse transcribed by transcriptase (Applied Biosystems, #4368813, Foster, CA). Transcript level was determined by SYBR Green gene expression assays (Applied Biosystems, #A25742). Real-time PCR reactions were carried out in triplicate using a qTOWER^3^G system (Analytik Jena). Primers were synthesized by IDT and their sequences are listed in Extended Data Table 2. The expression was calculated with comparative *Ct* method and the raw data was normalized by the internal control *Gapdh* or *18s*.

### Western blot

Cells were collected and lysed in RIPA buffer (Boston BioProducts, #BP-115) with proteinase inhibitor cocktail (Roche). Homogenized tissues were lysed using cell disruption buffer in Ambion® PARIS™ system. After 30 min incubation on ice and centrifugation, the supernatant was resolved on a 4-15% polyacrylamide gradient gel (BIO-RAD, #4568084 or # 5671085), transferred to 0.2 μm PVDF membrane (Thermo, #88520 or BIO-RAD, #1704157) and probed with corresponding antibodies. Antibody against CTSL was purchased from R&D SYSTEMS (#AF1515), against N protein from Sino Biological (#40143-R019, 1:1000), and against calnexin from Enzo (#ADI-SPA-860-F, 1:1000). Immunoblots were developed using SuperSignal West Pico PLUS chemiluminescent substrate (Thermo, # 34580) and visualized by LI-COR Odyssey Fc Imaging System. The protein expression was quantified by densitometry (ImageJ) and normalized to calnexin.

### Tissue virus titer by plaque assay

SARS-CoV-2 plaque assays were performed in the Duke Regional Biocontainment BSL3 Laboratory (Durham, NC) as previously described^47^. Serial dilutions of lung homogenate were incubated with Vero E6 cells in a standard plaque assay. Homogenate and cells were incubated at 37°C, 5% CO2 for 1 h. At the end of the incubation, 1 ml of a viscous overlay (1:1 2X DMEM and 1.2% methylcellulose) was added to each well. Plates were incubated for 4 days. After fixation, staining and washing, plates were dried and plaques from each dilution of sample were counted. Data are reported as plaque forming units per g of lung tissue (PFU/g).

### *In vitro* virus infection and assessment assays

#### Pseudotyped virus

Lentiviral pseudotyped virus was produced as previously described^48^. Briefly, HEK293T cells were transfected with HIV-1 NL4.3-inGluc (a gift of Marc Johnson at the University of Missouri, Columbia, Missouri, USA) and pcDNA3.1-SARS-CoV2-S-C9 (obtained from Fang Li at the University of Minnesota, St. Paul, Minnesota, USA) constructs in a 2:1 ratio using polyelthylenimine (PEI). Supernatants were harvested 24, 48, and 72 h post-transfection and were pooled, aliquoted, and stored at −80°C.

#### Pseudotyped virus infection

3 × 10^5^ of Vero-ACE2, Vero-ACE2-TMPRSS2 or Caco-2 were seeded into 6-well plate and transfected with *CasRx* mRNA and pre-gControl oligo (*CasRx*-pre-gControl) or pre-g*Ctsl* oligo (*CasRx*-pre-g*Ctsl*) by Lipofectamine MessengerMAX (Invitrogen, #LMRNA015). Cells were re-seeded into 24-well plate 12 h post-transfection and infected with pseudotyped virus for 6 h. The media was then replaced with 500 μl fresh media. 20 μl of media was collected at 24, 48, and 72 h and incubated with 20 μl of Gaussia luciferase substrate [0.1 M Tris (MilliporeSigma, #T6066) pH 7.4, 0.3 M sodium ascorbate (Spectrum, #S1349), 10 μM coelenterazine (GoldBio, #CZ2.5)] and read by BioTek Cytation5 plate-reader.

#### Authentic SARS-CoV-2 B.1.617.2 Delta variant and cell infection

The variant strain (USA/PHC658/2021) was obtained from BEI Resources NR-55611. 3 × 10^5^ of Vero-ACE2-TMPRSS2 cells were seeded into 6-well plate and transfected with *CasRx*-pre-gControl or *CasRx*-pre-g*Ctsl* by Lipofectamine MessengerMAX. Cells were re-seeded into 12-well plate 12 h post-transfection and infected with the SARS-Related Coronavirus 2, Isolate hCoV-19/USA/PHC658/2021 at MOI of 0.01.

#### TCID_50_ assay

Cell culture supernatants were collected 20 h after infection. Samples were serially diluted 10-fold and used to determine live virus titer by Tissue Culture Infection Dose 50 (TCID_50_). Briefly, 4 × 10^4^ Vero-E6 cells were placed in each well of a 96-well tissue culture plate and allowed to adhere overnight. Growth medium was removed, and serial dilutions of experimental supernatant were added. Samples in duplicate were incubated for 96 h, followed by being fixed with 10% NBF and stained with 0.1% crystal violet. Cytopathic effect was examined by eye, and TCID_50_/mL was calculated by the Reed-Muench method.

### Statistical analyses

All data were analyzed by GraphPad Prism 9.0 software. Differences in mean values between groups were determined by *Student*’s t-test or Mann-Whitney U-test as applicable. Differences in survival were analyzed by long-rank test. *P* values <0.05 were considered statistically significant. \**P*<0.05, \*\**P*<0.01, \*\*\**P*<0.001, \*\*\*\**P*<0.0001.

## Acknowledgements

We thank Amelia Karlsson, Kristina Riebe, Rose Asrican, Thomas Oguin III from the Duke Regional Biocontainment Laboratory (RBL) for performing authentic SARS-CoV-2 infection experiments *in vitro* and *vivo*. The RBL was partially constructed with funding from the NIH/NIAID (UC6-AI058607). We thank Zuowei Su from Duke BioRepository & Precision Pathology Center for performing IHC assay. We also thank Ling Wang from Animal Histopathology and Laboratory Medicine Core at UNC-Chapel Hill for performing animal hematological and clinical chemistry tests. We thank Marina Sokolsky from Nanomedicines Characterization Core Facility and Nanopformulation Core Facility at UNC-Chapel Hill for characterizing our LNPs. We thank Dr. Mark Wiesner (Duke) for allowing us to use his Zetasizer. The following reagents were deposited by the Centers for Disease Control and Prevention and obtained through BEI Resources, NIAID, NIH: SARS-Related Coronavirus 2, Isolate USA-WA1/2020, NR-52281; SARS-Related Coronavirus 2, Isolate hCoV-19/USA/PHC658/2021 (Lineage B.1.617.2; Delta Variant), NR-55611, contributed by Dr. Richard Webby and Dr. Anami Patel.

## Author contributions

Conceived project: Q.W; Conceived and designed study: QW, SLL, YD, HW, ZC; Designed and performed experiments: ZC, CZ, FH, FY, JY, YZ, GDS; Data analysis: ZC, CZ, JE; Manuscript writing: ZC, HW, JH, HS, JE, GDS, YD, SLL, QW.

## Competing interests

QW, ZC, and YD are inventors on a patent filed by Duke University that relates to the research reported in this paper. JH is a consultant for or owns shares in the following companies: Kingmed, MoreHealth, OptraScan, Genetron, Omnitura, Vetonco, York Biotechnology, Genecode, VIVA Biotech and Sisu Pharma, and received grants from Zenith Epigenetics, BioXcel Therapeutics, Inc., and Fortis Therapeutics.

---

**Extended Data Fig. 1.**
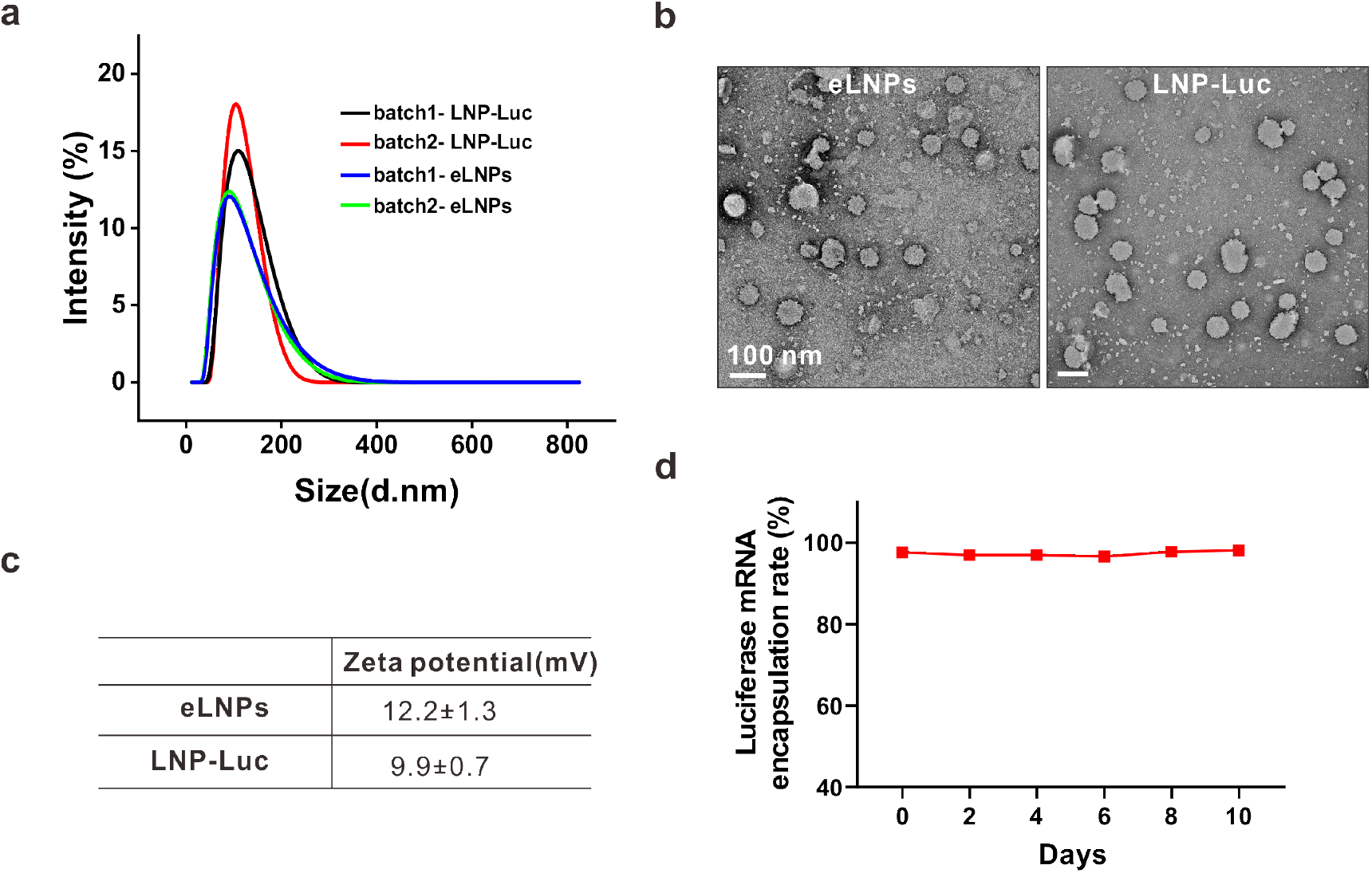
Characterization of empty LNPs (eLNPs) and LNPs encapsulating luciferase mRNA (LNP-Luc). **a**, Size distribution. The size of two batches of eLNPs and LNP-Luc were analyzed by Zetasizer. **b**, Representative transmission electron microscopic images. **c**, Zeta potential. Data are presented as mean ± SD of two batches of samples. **d**, Encapsulation rate and stability of LNP-Luc. The encapsulation efficiency of LNP-Luc was determined by Ribogreen assay on Day 0, 2, 4, 6, 8,10 days following its storage at 4°C.

**Extended Data Fig. 2.**
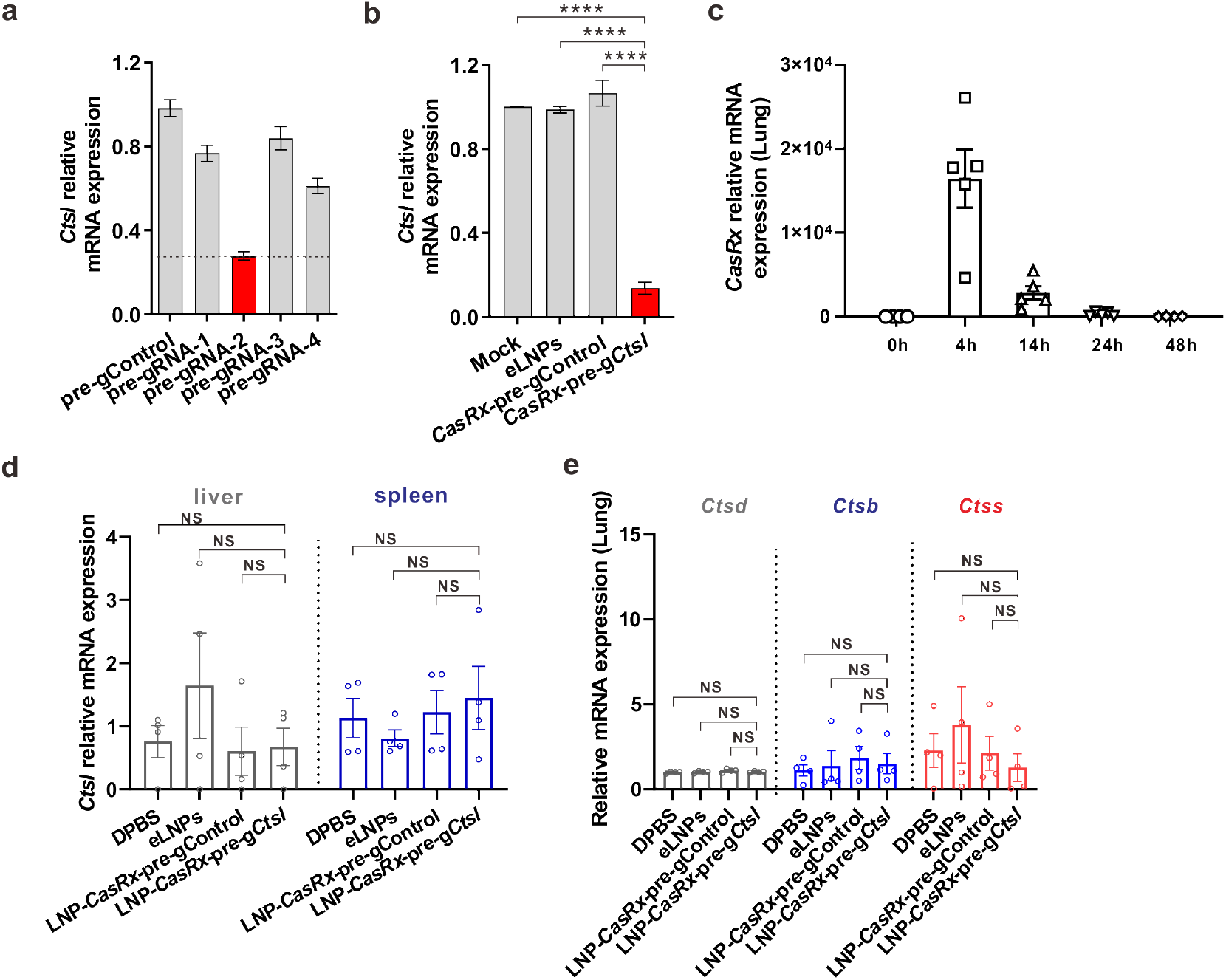
Screening for pre-gRNA targeting mouse *Ctsl in vitro* and examination for the editing specificity of LNP-*CasRx*-pre-g*Ctsl in vivo*. Relative *Ctsl* mRNA expression was measured by RT-PCR after 48 h transfection with **a**, Plasmids of CasRx with different pre-gRNAs designed for targeting of mouse *Ctsl*; or **b**, *CasRx* mRNA and selected pre-g*Ctsl* from (**a**) in mouse KLN205 cells. The experiments were repeated with each performed in triplicates. Data are presented as mean ± SEM. *P* values were calculated by two-tailed *Student*’s t-tests. **** *P*<0.0001. **c,** Kinetics of *CasRx* mRNA in lung. C57BL/6 mice were given LNPs encapsulating *CasRx* mRNA at 0.5 mg/kg by tail vein. Lungs were collected at 0, 4, 14, 24 and 48 h, respectively, after the injection for qRT-PCR analysis. Data are presented as mean ± SEM for each time point (n=4 for 0, 48 h; n=5 for 4 h, 14 h, 24 h). **d-e,** C57BL/6 mice were given DPBS, eLNPs, LNP-*CasRx*-pre-gControl or LNP-*CasRx*-pre-g*Ctsl* by tail vein. After 72 h, the lung, liver and spleen were collected for analysis. **d**, *Ctsl* mRNA levels in liver and spleen are unchanged by LNP-*CasRx*-pre-g*Ctsl* treatment. **e**, mRNA levels of isoform *Ctsd, Ctsb* and *Ctss* in lung are unchanged by LNP-*CasRx*-pre-g*Ctsl* treatment. Transcript levels are normalized by *Gapdh* and presented as mean ± SEM. n=4 per group, 2 females and 2 males. *P* values were calculated by one-tailed Mann-Whitney u test. NS, not significant.

**Extended Data Fig. 3.**
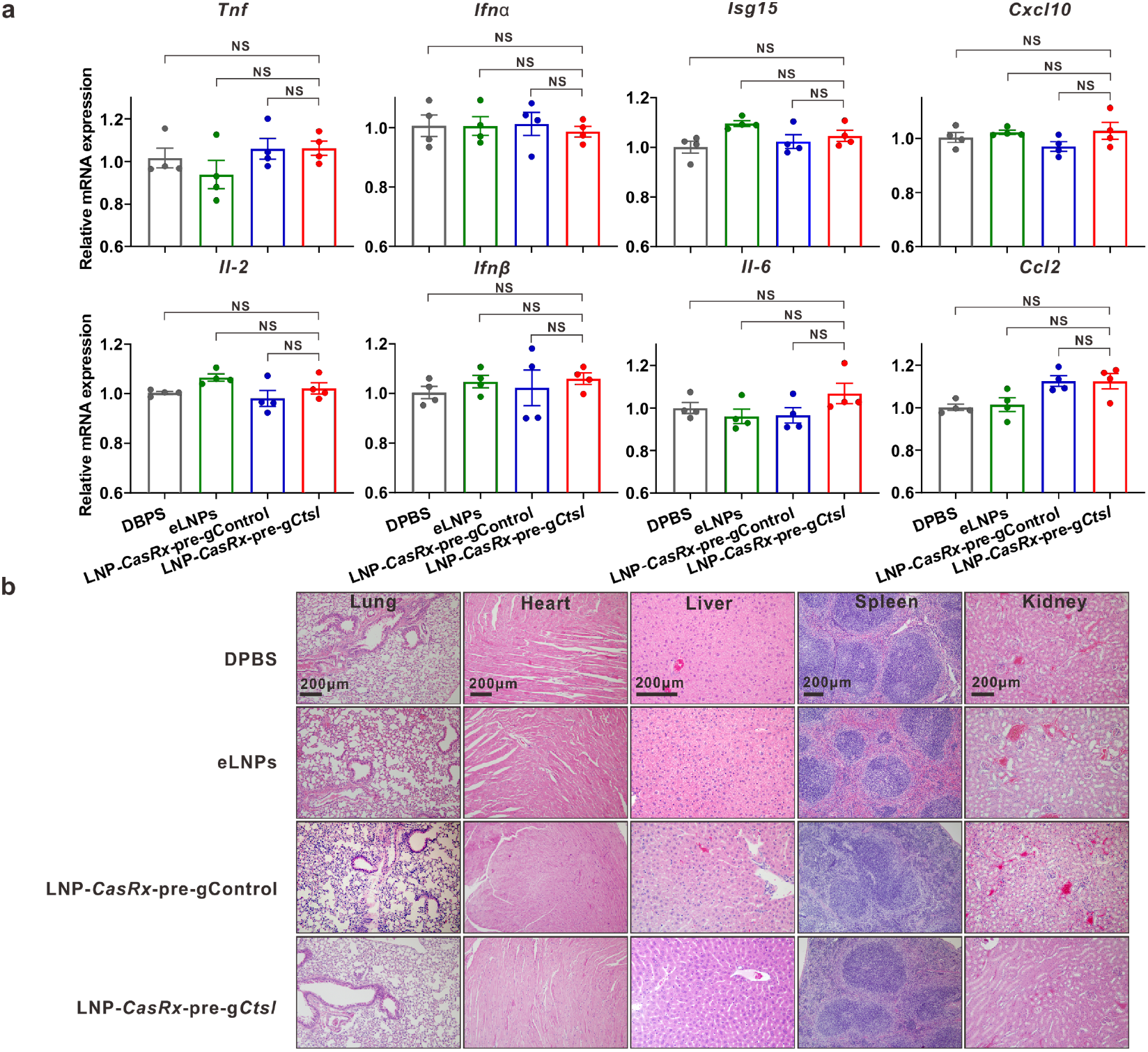
Immunogenicity and histological evaluation of LNP-*CasRx*-pre-g*Cts1* treatment in C57BL/6 mice. **a**, Cytokine and chemokine transcript levels in mouse lung are unchanged by LNP-*CasRx*-pre-g*Ctsl* treatment. Tissues were collected 72 h following the treatments. All transcript levels are normalized by *Gapdh*. Data are presented as mean ± SEM for each group (n=4, 2 females and 2 males). *P* values were calculated by two-tailed Mann-Whitney u test, NS, not significant. **b**, No substantial histopathological changes produced by LNP-*CasRx*-pre-g*Ctsl* treatment in indicated organ tissues. Two sections of each organ from 3 mice per group were evaluated after H&E staining and the representative images are presented.

**Extended Data Fig. 4.**
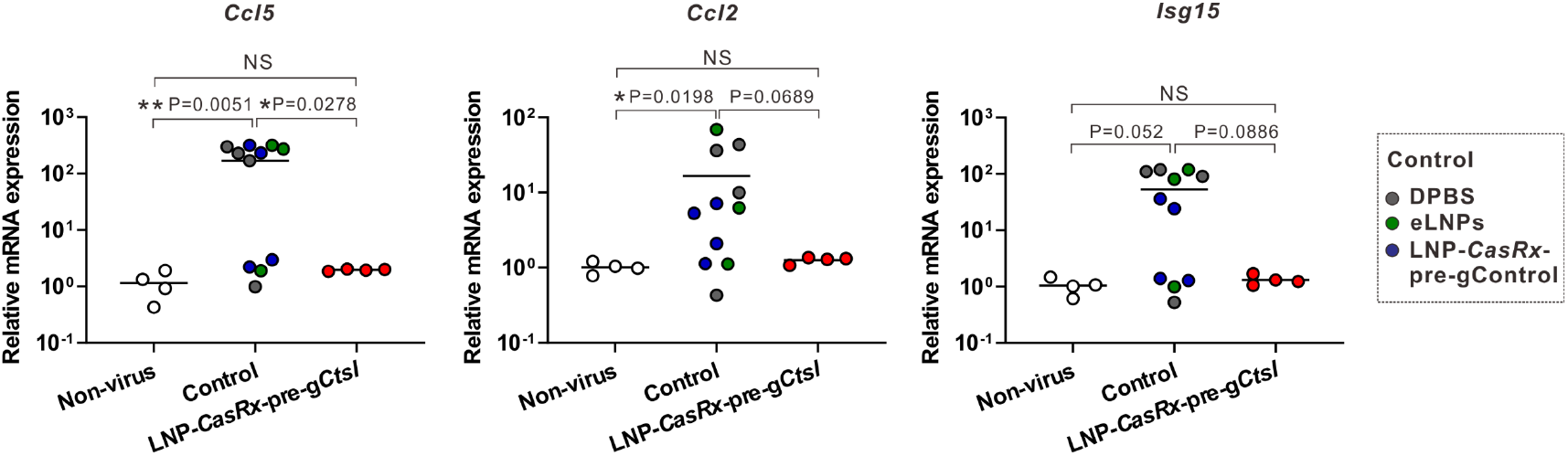
CasRx-mediated Ctsl knockdown reduces chemokines/cytokines in lungs of K18-hACE2 mice on Day 4 after SARS-CoV-2 infection. The experiment design is same as in Fig. 4. *Ccl5, Ccl2* and *Isg15* transcript levels in lung were determined by qRT-PCR and normalized by *Gapdh. P* values were calculated by one-tailed Mann-Whitney u test, grand mean. NS, not significant, * *P*<0.05, ** *P*<0.01.

**Extended Data Fig. 5.**
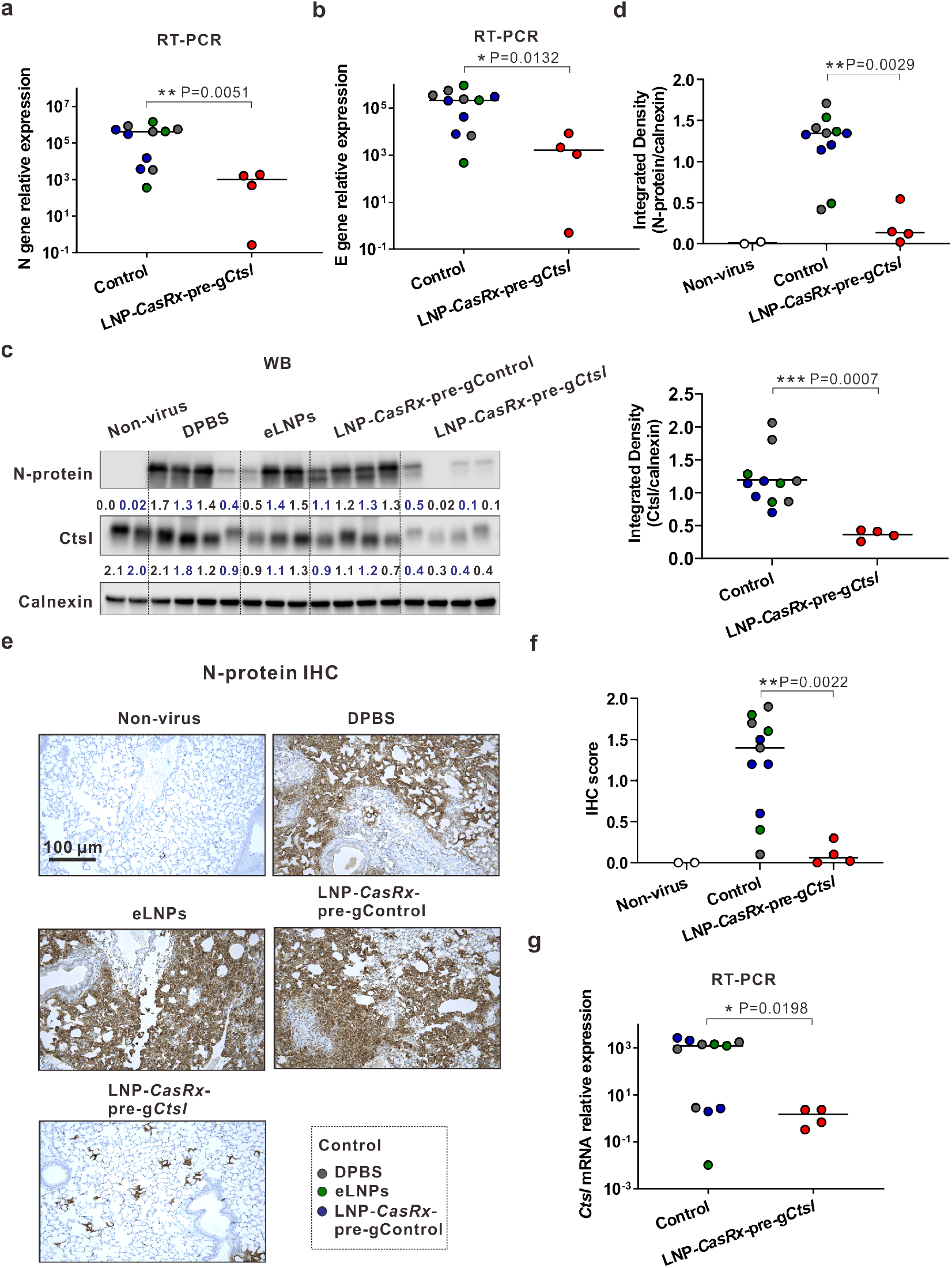
CasRx-mediated Ctsl knockdown reduces virus burden in lungs of K18-hACE2 mice on Day 4 after SARS-CoV-2 infection. **a**, Viral N gene transcript level. **b**, Viral E gene transcript level. **c**, Western blot of viral N protein and Ctsl in mouse lung. Calnexin was used as a loading control. The ratio of N protein or Ctsl over calnexin is listed under the blot. Non-virus, two mice (1 female and 1 male) were not challenged with virus. **d**, Integrated density of N protein (upper panel) and Ctsl (down panel) over the loading control. **e**, Representative images for N protein immunostaining in lung. **f**, IHC score for N protein. **g**, *Ctsl* transcript levels. **a**, **b, g**, All transcript levels are normalized by *Gapdh. P* values were calculated by one-tailed Mann-Whitney u test, grand mean. * *P*<0.05, ** *P*<0.01, *** *P*<0.001.

**Extended Data Fig. 6.**
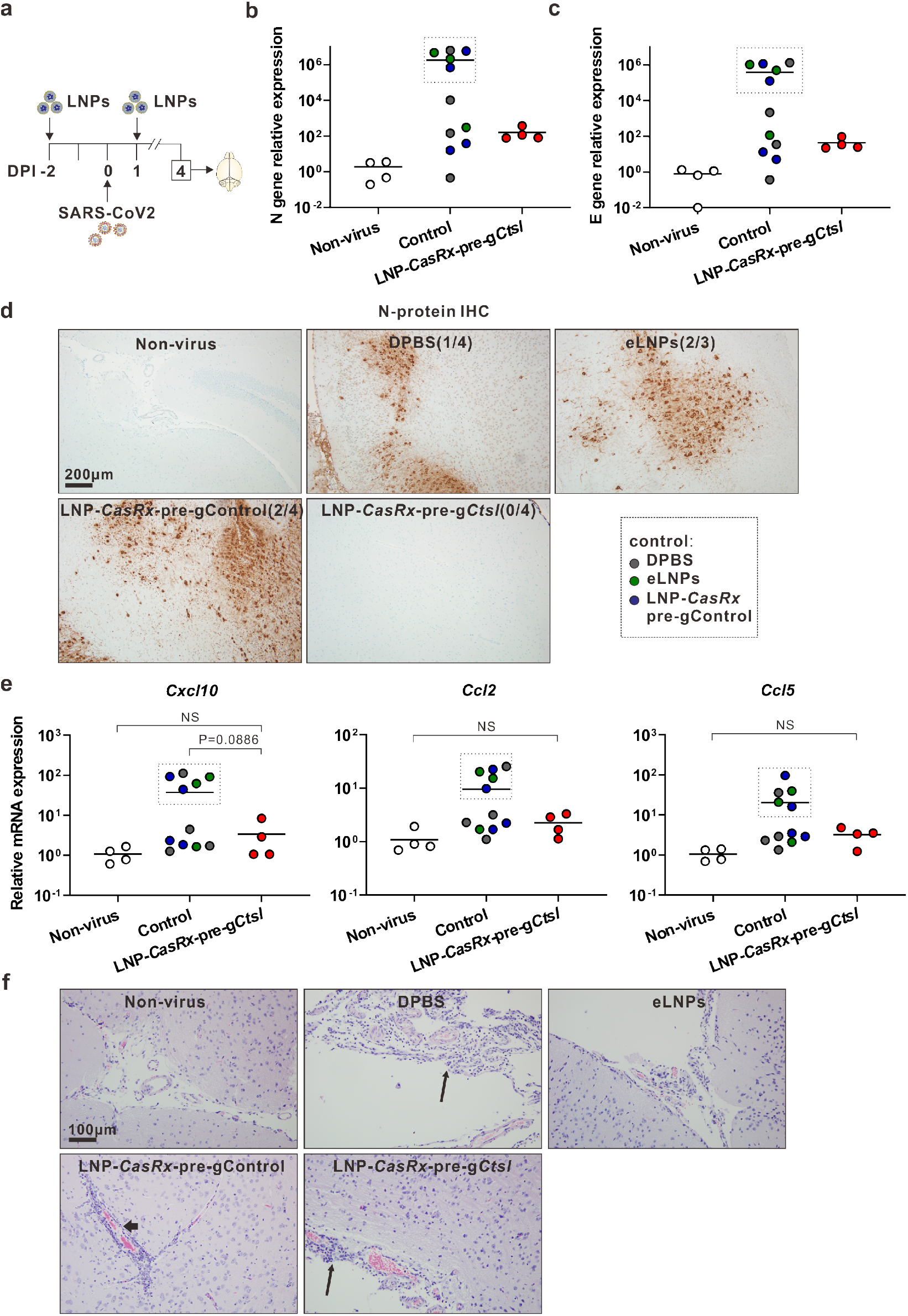
Brain responses in SARS-CoV-2-infected K18-hACE2 mice at 4 DPI following LNP-*CasRx*-pre-g*Ctsl* treatment. **a**, Schematic illustration of the experiment design. The treatments and virus challenge are same as in Fig. 2. Brains were collected at 4 DPI for analysis. The right lobes were subjected for RT-PCR, whereas the sections of the left lobes for IHC staining. **b**, Viral N gene transcript level. **c**, Viral E gene transcript level. **d**, Representative photomicrographs of N-protein immunostaining shows strong intracytoplasmic immunoreactivity of neurons in cerebral cortex, thalamus, and hypothalamus regions. Two sections of brain tissues from 4 mice of each group (3 for eLNPs group) were subjected to IHC analysis. The occurrence of strong positive staining was indicated in brackets for each group. **e**, *Cxcl10, Ccl2*, and *Ccl5* transcript levels. **f**, Representative photomicrographs of perivascular lymphoid infiltrate (thin arrows) in meninges and Virchow-Robin spaces (thick arrow) demonstrates vascular inflammatory changes in all groups. **b, c, e**, all transcript levels were normalized by *Gapdh. P* values were calculated by one-tailed Mann-Whitney u test, grand mean. NS, not significant. As boxed in the graphs, 5 out of 11 mice in the Control group express high transcript levels of viral N gene, E gene and chemokines in brain.

**Extended Data Fig. 7.**
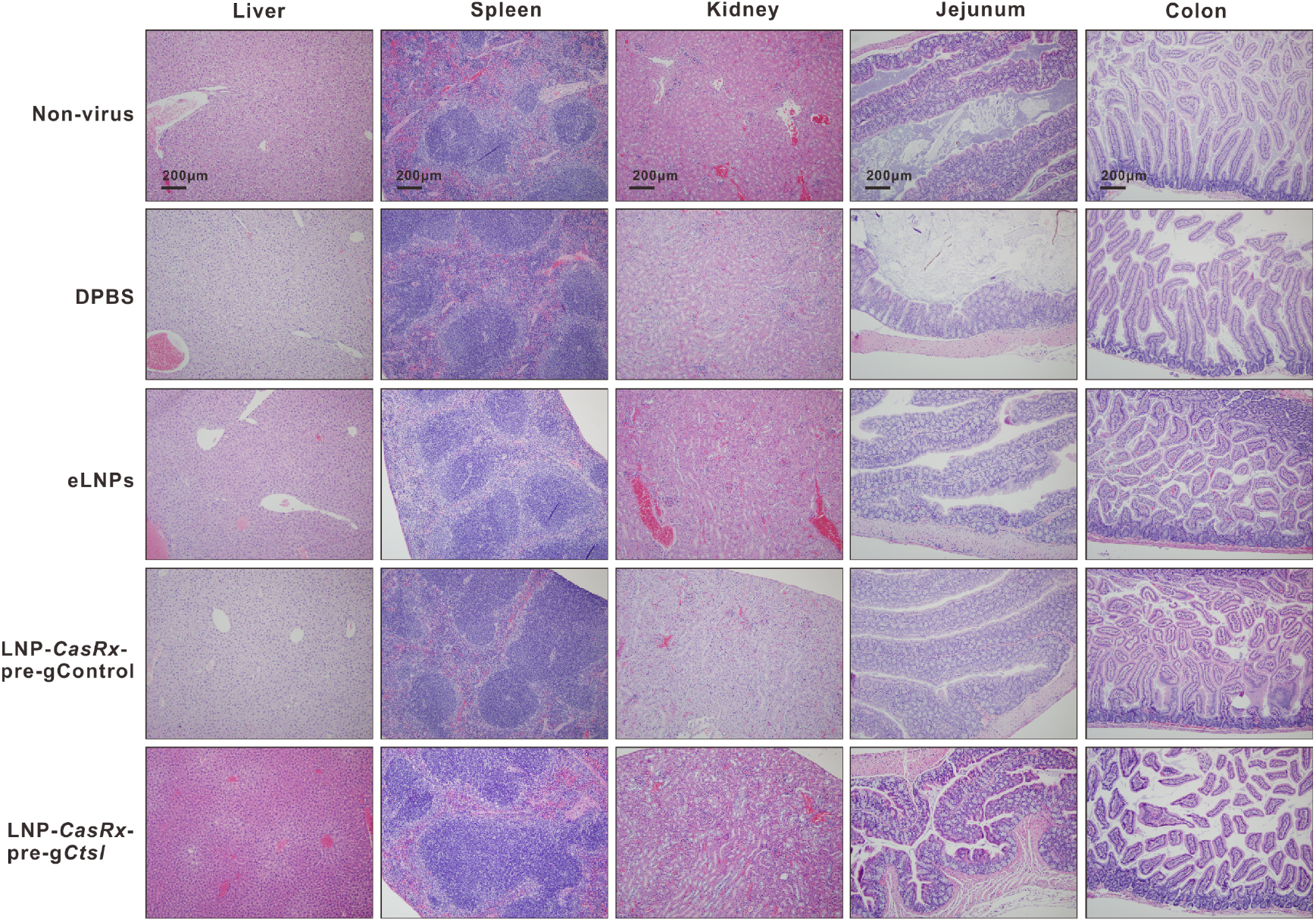
No histopathological changes observed in organs of liver, spleen, kidney, jejunum, and colon in SARS-CoV-2-infected K18-hACE2 mice at 4 DPI following LNP-*CasRx*-pre-g*Ctsl* treatment.

**Extended Data Fig. 8.**
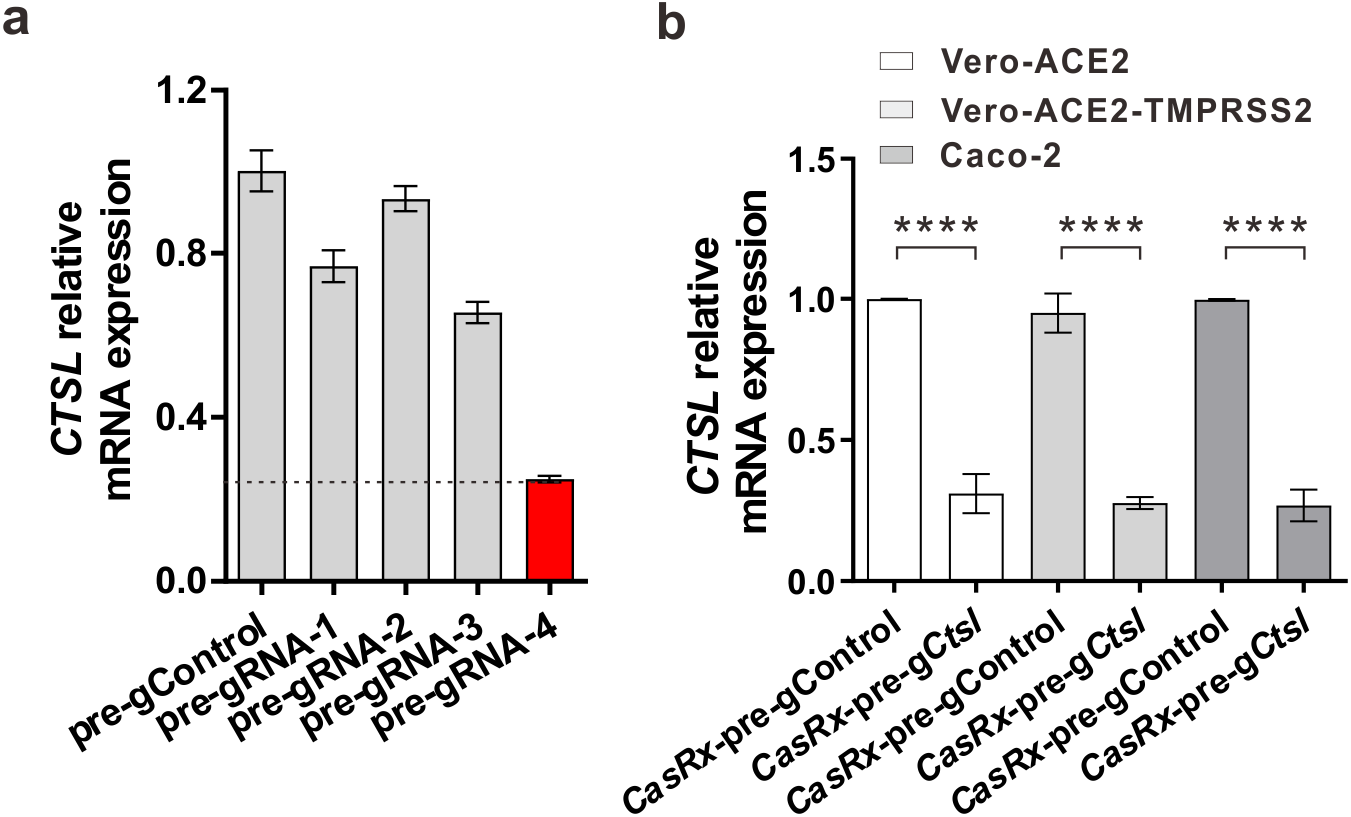
Screening for pre-gRNA targeting human *CTSL* and evaluation of its knockdown efficiency in cultured cells. **a,** Relative *CTSL* mRNA expression was measured by RT-PCR after 48 h transfection with CasRx plasmid and different pre-gRNAs designed for targeting of human *CTSL* in 293 FT cells. **b,** *CTSL* knockdown efficiency in Vero-ACE2, Vero-ACE2-TMPRSS2 and Caco-2 cells. The experiment was repeated three times with each performed in triplicate. Transcript levels are normalized by 18s and presented as mean ± SEM. *P* values were calculated by *Student*’s t-test. **** *P*<0.0001.

## References

1 Hoffmann, M. et al. SARS-CoV-2 Cell Entry Depends on ACE2 and TMPRSS2 and Is Blocked by a Clinically Proven Protease Inhibitor. Cell 181, 271–280 e278, doi:10.1016/j.cell.2020.02.052 (2020).

2 Wei, J. et al. Genome-wide CRISPR Screens Reveal Host Factors Critical for SARS-CoV-2 Infection. Cell 184, 76–91 e13, doi:10.1016/j.cell.2020.10.028 (2021).

3 Daniloski, Z. et al. Identification of Required Host Factors for SARS-CoV-2 Infection in Human Cells. Cell 184, 92–105 e116, doi:10.1016/j.cell.2020.10.030 (2021).

4 Beigel, J. H. et al. Remdesivir for the Treatment of Covid-19 - Final Report. N Engl J Med 383, 1813–1826, doi:10.1056/NEJMoa2007764 (2020).

5 Rubin, D., Chan-Tack, K., Farley, J. & Sherwat, A. FDA Approval of Remdesivir - A Step in the Right Direction. N Engl J Med 383, 2598–2600, doi:10.1056/NEJMp2032369 (2020).

6 Taylor, P. C. et al. Neutralizing monoclonal antibodies for treatment of COVID-19. Nat Rev Immunol 21, 382–393, doi:10.1038/s41577-021-00542-x (2021).

7 Marovich, M., Mascola, J. R. & Cohen, M. S. Monoclonal Antibodies for Prevention and Treatment of COVID-19. JAMA 324, 131–132, doi:10.1001/jama.2020.10245 (2020).

8 Mlcochova, P. et al. SARS-CoV-2 B.1.617.2 Delta variant replication and immune evasion. Nature, doi:10.1038/s41586-021-03944-y (2021).

9 Gordon, D. E. et al. A SARS-CoV-2 protein interaction map reveals targets for drug repurposing. Nature 583, 459–468, doi:10.1038/s41586-020-2286-9 (2020).

10 Liu, T., Luo, S., Libby, P. & Shi, G. P. Cathepsin L-selective inhibitors: A potentially promising treatment for COVID-19 patients. Pharmacol Ther 213, 107587, doi:10.1016/j.pharmthera.2020.107587 (2020).

11 Ou, X. et al. Characterization of spike glycoprotein of SARS-CoV-2 on virus entry and its immune cross-reactivity with SARS-CoV. Nat Commun 11, 1620, doi:10.1038/s41467-020-15562-9 (2020).

12 Smieszek, S. P., Przychodzen, B. P. & Polymeropoulos, M. H. Amantadine disrupts lysosomal gene expression: A hypothesis for COVID19 treatment. Int J Antimicrob Agents 55, 106004, doi:10.1016/j.ijantimicag.2020.106004 (2020).

13 Zhou, Y. et al. Protease inhibitors targeting coronavirus and filovirus entry. Antiviral Res 116, 76–84, doi:10.1016/j.antiviral.2015.01.011 (2015).

14 Simmons, G. et al. Inhibitors of cathepsin L prevent severe acute respiratory syndrome coronavirus entry. Proc Natl Acad Sci U S A 102, 11876–11881, doi:10.1073/pnas.0505577102 (2005).

15 Zhao, M. M. et al. Cathepsin L plays a key role in SARS-CoV-2 infection in humans and humanized mice and is a promising target for new drug development. Signal Transduct Target Ther 6, 134, doi:10.1038/s41392-021-00558-8 (2021).

16 Steuten, K. et al. Challenges for Targeting SARS-CoV-2 Proteases as a Therapeutic Strategy for COVID-19. ACS Infect Dis 7, 1457–1468, doi:10.1021/acsinfecdis.0c00815 (2021).

17 Cheng, Q. et al. Selective organ targeting (SORT) nanoparticles for tissue-specific mRNA delivery and CRISPR-Cas gene editing. Nat Nanotechnol 15, 313–320, doi:10.1038/s41565-020-0669-6 (2020).

18 Mullard, A. FDA approves landmark RNAi drug. Nat Rev Drug Discov 17, 613, doi:10.1038/nrd.2018.152 (2018).

19 Konermann, S. et al. Transcriptome Engineering with RNA-Targeting Type VI-D CRISPR Effectors. Cell 173, 665–676 e614, doi:10.1016/j.cell.2018.02.033 (2018).

20 Eoh, J. & Gu, L. Biomaterials as vectors for the delivery of CRISPR-Cas9. Biomater Sci 7, 1240–1261, doi:10.1039/c8bm01310a (2019).

21 Tong, S., Moyo, B., Lee, C. M., Leong, K. & Bao, G. Engineered materials for in vivo delivery of genome-editing machinery. Nat Rev Mater 4, 726–737 (2019).

22 Yin, H. et al. Therapeutic genome editing by combined viral and non-viral delivery of CRISPR system components in vivo. Nat Biotechnol 34, 328–333, doi:10.1038/nbt.3471 (2016).

23 Oladunni, F. S. et al. Lethality of SARS-CoV-2 infection in K18 human angiotensin-converting enzyme 2 transgenic mice. Nat Commun 11, 6122, doi:10.1038/s41467-020-19891-7 (2020).

24 Rathnasinghe, R. et al. Comparison of transgenic and adenovirus hACE2 mouse models for SARS-CoV-2 infection. Emerg Microbes Infect 9, 2433–2445, doi:10.1080/22221751.2020.1838955 (2020).

25 Zheng, J. et al. COVID-19 treatments and pathogenesis including anosmia in K18-hACE2 mice. Nature 589, 603–607, doi:10.1038/s41586-020-2943-z (2021).

26 Yinda, C. K. et al. K18-hACE2 mice develop respiratory disease resembling severe COVID-19. PLoS Pathog 17, e1009195, doi:10.1371/journal.ppat.1009195 (2021).

27 Munoz-Fontela, C. et al. Animal models for COVID-19. Nature 586, 509–515, doi:10.1038/s41586-020-2787-6 (2020).

28 Winkler, E. S. et al. SARS-CoV-2 infection of human ACE2-transgenic mice causes severe lung inflammation and impaired function. Nat Immunol 21, 1327–1335, doi:10.1038/s41590-020-0778-2 (2020).

29 Coperchini, F., Chiovato, L., Croce, L., Magri, F. & Rotondi, M. The cytokine storm in COVID-19: An overview of the involvement of the chemokine/chemokine-receptor system. Cytokine Growth Factor Rev 53, 25–32, doi:10.1016/j.cytogfr.2020.05.003 (2020).

30 Song, P., Li, W., Xie, J., Hou, Y. & You, C. Cytokine storm induced by SARS-CoV-2. Clin Chim Acta 509, 280–287, doi:10.1016/j.cca.2020.06.017 (2020).

31 Del Valle, D. M. et al. An inflammatory cytokine signature predicts COVID-19 severity and survival. Nat Med 26, 1636–1643, doi:10.1038/s41591-020-1051-9 (2020).

32 Chou, S. H. et al. Global Incidence of Neurological Manifestations Among Patients Hospitalized With COVID-19-A Report for the GCS-NeuroCOVID Consortium and the ENERGY Consortium. JAMA Netw Open 4, e2112131, doi:10.1001/jamanetworkopen.2021.12131 (2021).

33 Liotta, E. M. et al. Frequent neurologic manifestations and encephalopathy-associated morbidity in Covid-19 patients. Ann Clin Transl Neurol 7, 2221–2230, doi:10.1002/acn3.51210 (2020).

34 Romero-Sanchez, C. M. et al. Neurologic manifestations in hospitalized patients with COVID-19: The ALBACOVID registry. Neurology 95, e1060–e1070, doi:10.1212/WNL.0000000000009937 (2020).

35 Yang, A. C. et al. Dysregulation of brain and choroid plexus cell types in severe COVID-19. Nature 595, 565–571, doi:10.1038/s41586-021-03710-0 (2021).

36 Lopez-Otin, C. & Bond, J. S. Proteases: multifunctional enzymes in life and disease. J Biol Chem 283, 30433–30437, doi:10.1074/jbc.R800035200 (2008).

37 Yadati, T., Houben, T., Bitorina, A. & Shiri-Sverdlov, R. The Ins and Outs of Cathepsins: Physiological Function and Role in Disease Management. Cells 9, doi:10.3390/cells9071679 (2020).

38 Birmingham, A. et al. 3’ UTR seed matches, but not overall identity, are associated with RNAi off-targets. Nat Methods 3, 199–204, doi:nmeth854 [pii] 10.1038/nmeth854 (2006).

39 Sigoillot, F. D. et al. A bioinformatics method identifies prominent off-targeted transcripts in RNAi screens. Nat Methods 9, 363–366, doi:10.1038/nmeth.1898 (2012).

40 Beers, C. et al. Cathepsin S controls MHC class II-mediated antigen presentation by epithelial cells in vivo. J Immunol 174, 1205–1212, doi:10.4049/jimmunol.174.3.1205 (2005).

41 Shi, G. P. et al. Cathepsin S required for normal MHC class II peptide loading and germinal center development. Immunity 10, 197–206, doi:10.1016/s1074-7613(00)80020-5 (1999).

42 Honey, K., Nakagawa, T., Peters, C. & Rudensky, A. Cathepsin L regulates CD4+ T cell selection independently of its effect on invariant chain: a role in the generation of positively selecting peptide ligands. J Exp Med 195, 1349–1358, doi:10.1084/jem.20011904 (2002).

43 Nakagawa, T. et al. Cathepsin L: critical role in Ii degradation and CD4 T cell selection in the thymus. Science 280, 450–453, doi:10.1126/science.280.5362.450 (1998).

44 Meganck, R. M. & Baric, R. S. Developing therapeutic approaches for twenty-first-century emerging infectious viral diseases. Nat Med 27, 401–410, doi:10.1038/s41591-021-01282-0 (2021).

45 Li, B., Luo, X. & Dong, Y. Effects of Chemically Modified Messenger RNA on Protein Expression. Bioconjug Chem 27, 849–853, doi:10.1021/acs.bioconjchem.6b00090 (2016).

46 Li, B., Zeng, C. & Dong, Y. Design and assessment of engineered CRISPR-Cpf1 and its use for genome editing. Nat Protoc 13, 899–914, doi:10.1038/nprot.2018.004 (2018).

47 Saunders, K. O. et al. Neutralizing antibody vaccine for pandemic and pre-emergent coronaviruses. Nature 594, 553–559, doi:10.1038/s41586-021-03594-0 (2021).

48 Zeng, C. et al. Neutralizing antibody against SARS-CoV-2 spike in COVID-19 patients, health care workers, and convalescent plasma donors. JCI Insight 5, doi:10.1172/jci.insight.143213 (2020).

